# Dissimilatory Nitrate Reduction to Ammonium in the Cable Bacterium *Ca*. Electronema Sp. GS

**DOI:** 10.1101/2020.02.22.960823

**Authors:** Ugo Marzocchi, Casper Thorup, Ann-Sofie Dam, Andreas Schramm, Nils Risgaard-Petersen

**Author notes:** Corresponding author Address: Ugo Marzocchi, Center for Electromicrobiology, Department of Bioscience at Aarhus University. Ny Munkegade 114, 8000-C Aarhus, Denmark.

## Abstract

Cable bacteria are filamentous Desulfobulbaceae that split the energy-conserving reaction of sulphide oxidation into two half reactions occurring in distinct cells. Cable bacteria can use nitrate, but the reduction pathway is unknown, making it difficult to assess their direct impact on the N-cycle. Here we show that the freshwater cable bacterium *Ca.* Electronema sp. GS performs dissimilatory nitrate reduction to ammonium (DNRA). ^15^NO_3_^−^-amended sediment with *Ca.* Electronema sp. GS showed higher rates of DNRA and nitrite production than sediment without *Ca.* Electronema sp. GS. Electron flux from sulphide oxidation, inferred from electric potential measurements, matched the electron flux needed to drive cable bacteria-mediated nitrate reduction. *Ca.* Electronema sp. GS expressed a complete *nap* operon for periplasmic nitrate reduction to nitrite, and genes encoding a periplasmic multiheme cytochrome (pMHC), homolog to a pMHC that can catalyse nitrite reduction to ammonium in *Ca.* Maribeggiatoa. Phylogenetic analysis suggests that the capacity for DNRA was acquired in multiple events through horizontal gene transfer from different organisms, before cable bacteria split into different salinity niches. The architecture of the nitrate reduction system suggests absence of energy conservation through oxidative phosphorylation, indicating that cable bacteria primarily conserve energy through the half reaction of sulfide oxidation.

## INTRODUCTION

Cable bacteria (CB) are centimetre-long filamentous Desulfobulbaceae. They are able to split the energy-conserving reaction of sulphide oxidation into two half-redox reactions: anodic sulphide oxidation and cathodic oxygen reduction and distribute these among distinct cells (1). This is possible, as all cells in the filament are interlinked by highly conductive strings (1, 2), so that cells with access to sulphide can perform anodic sulphide oxidation donating electrons to the strings, whereas cells with access to oxygen can accept electrons from the strings and perform cathodic oxygen reduction. The overall metabolic process was first discovered in marine sediments (3) and is named electrogenic sulphide oxidation (e-SOx) (4). Since their discovery, cable bacteria and e-SOx have been found in diverse aquatic environments, such as marine, brackish, and freshwater sediments (5–7), where they can significantly influence the cycling of major elements such as sulphur, iron, oxygen, and carbon (8–10).

In the absence of oxygen, CB can use nitrate or nitrite as electron acceptor in e-SOx (11, 12). Experiments conducted on riverine sediment show that e-SOx can promote sediment DNRA activity indirectly, via increasing the Fe^2+^ pool through dissolution of FeS minerals (13). However, as the exact nitrate reduction pathway by CB remains unknown, it is at present not possible to fully understand the direct and therefore overall impact of cable bacteria on the cycling of nitrogen. Recent CB genome sequencing reported the presence of genes encoding the periplasmic reductase NapAB that catalyses the reduction of nitrate to nitrite (14). From incubation experiments it can be deduced that marine CB can reduce nitrite but also that they lack a full denitrification pathway, as these experiments showed that nitrous oxide could not act as electron acceptor (12). Typical, *nir* or *nrf*-type, nitrite reductases are absent in CB genomes, but a periplasmic multi-haem cytochrome (pMHC) homologous to a pMHC previously found in the orange sulphur bacteria *Ca.* Maribeggiatoa sp. is encoded in the most complete CB genomes (14). The pMHC in *Ca.* Maribeggiatoa sp. has been shown to catalyse the reduction of nitrite to ammonium *in vitro* (15). Hence a likely pathway for nitrate reduction in CB is nitrate reduction to nitrite followed by nitrite reduction to ammonium, *i.e.* the dissimilatory nitrate reduction to ammonium (DNRA).

In the present study, we used a combined ^15^N and transcriptome approach to test the hypothesis that cable bacteria perform DNRA. The model organism in our study was an enrichment of the freshwater CB strain *Ca.* Electronema sp. GS. This strain can be cultivated in autoclaved sediment and its genome has been recently described (14).

## MATERIALS AND METHODS

### *Ca.* Electronema sp. GS sediment enrichment and incubation

Sediment collected from a freshwater lake (Vennelyst Sø, Aarhus, Denmark) was sieved trough a 0.5 mm mesh, homogenized, and autoclaved. The autoclaved sediment was packed into eight Plexiglas® liners (inner diam. 4.3 cm, Fig. 1A and B). The top layer of seven cores was inoculated with a few grams of a sediment enrichment culture of *Ca.* Electronema sp. GS (14). These are in the following assigned as “CB-cores”. Another four cores were left un-inoculated. These are in the following assigned as “CB-free cores”. The CB- and CB-free cores were then pre-incubated for four weeks in two separate aquaria with tap water, aerated with submerged air pumps and held at 15^°^C, to promote the growth of *Ca.* Electronema sp. GS in the CB-cores, as CB have been shown to grow faster in the presence of oxygen than under anoxic conditions with nitrate (11). The water in both aquaria was replaced weekly to replenish nutrients. The development of an active *Ca.* Electronema sp. GS population was monitored through measurements of the electric potential (EP) distribution in the cores and through regular microscopic inspections of the sediment.

**Figure 1.**
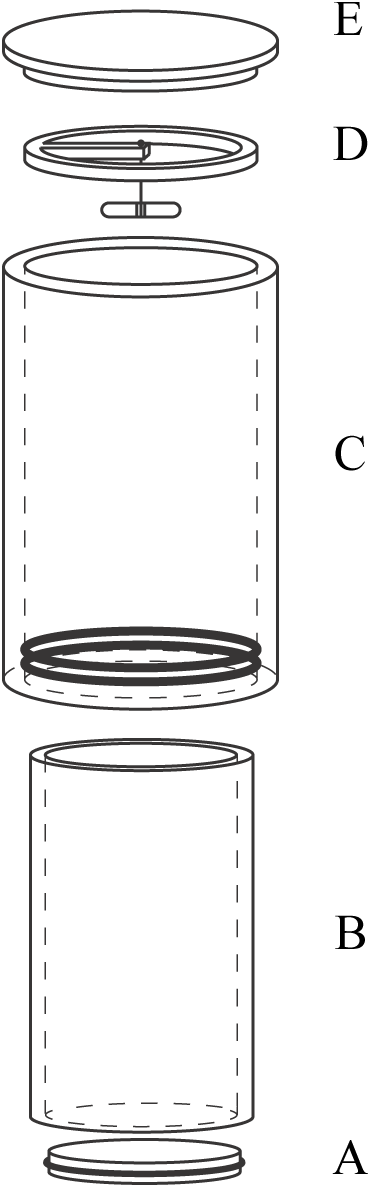
Core liner and water enclosure for sediment incubation. Bottom stopper with o-ring (A), liner for sediment incubation (B), enclosure for water incubation with o-rings (C), magnetic stirring bar and holder (D), top lid (E).

Following the pre-incubation period, three CB-cores were sampled for RNA extraction and served as oxic controls. Then, both aquaria were sealed and the water purged with pre-mixed N_2_ and CO_2_ (0.04 %) gas to remove dissolved oxygen while maintaining constant pH. The oxygen concentration was <0.2% air saturation as monitored with fiber-optic O_2_ sensors (FireStingO2, Pyroscience, Germany). A 1:1 mixture of K^14^NO_3_ and K^15^NO_3_ (^15^N-atom%: 98%; Sigma-Aldrich) was added to a final concentration of 400 µM. The nitrate concentration was kept constant by adding small aliquots from a 100 mM K^14^NO_3_: K^15^NO_3_ stock solution two to three times per week. After two days of exposure to anoxia and nitrate, the activity of the *Ca.* Electronema sp. GS population was assessed through measurements of the EP distribution in the CB-cores. After three days of exposure, N-fluxes at the sediment-water interface were measured over a period of ten days to assess the rates of nitrate reduction in the CB- and the CB-free cores. Thereafter, sediment samples for RNA analysis where collected from the CB-cores.

### The electric potential distribution and e-SOx activity

The activity of the *Ca.* Electronema sp. GS. population during the pre-incubation period and during nitrate exposure was inferred from the depth distribution of the EP as described previously (12). Depth distribution of the EP was measured with a customized EP microelectrode (16) against a Red Rod reference electrode (REF201 Radiometer Analytical, Denmark) kept in the overlying water. Both electrodes were connected to an in-house-made milli-voltmeter with a resistance >10^14^ Ω (Aarhus University, Denmark). The analog signal from the milli-voltmeter was digitized for PC-processing using a 16-bit A/D converter (ADC-216; Unisense A/S, Denmark). To control its stepwise movement, the microelectrode was fixed to a 1-D microprofiling system (Unisense A/S, Denmark). The software Sensor Trace PRO was used to control the movements of the sensor and for logging the sensor signals. During the pre-incubation period, the EP-distribution was measured four times in the CB- and twice in the CB-free cores to evaluate the establishment (or lack) of e-SOx activity.

During the nitrate exposure period, the EP-distribution was measured only in the CB-cores to quantify the overall e-SOx activity of the *Ca.* Electronema sp. GS population and its ability to perform cathodic nitrate reduction. For this, cores were removed from the aquarium and fitted with a top Plexiglass® tube (Fig. 1C, no lid on) that allowed to maintain an overlying water column of 150 mL. The tube was filled with nitrate-free water kept aerated (100% air saturation) and gently stirred with an air pump. Cable bacteria activity was tested by sediment EP microprofiling as described above. Successively, to evaluate the magnitude of the “residual” EP (*i.e.*, not due to e-SOx activity), EP profiling was repeated after oxygen was removed from the water by gentle purging with pre-mixed N_2_ and CO_2_ (0.04 %) gas. Finally, to test the ability of *Ca.* Electronema sp. GS to perform cathodic nitrate reduction, EP profiling was conducted after the addition of a concentrated (10 mM) KNO_3_ anoxic solution to the water (final conc.: 200 µM). The magnitude of the electron flux J was calculated for the CB-cores exposed to oxic or anoxic water with nitrate according to (12):

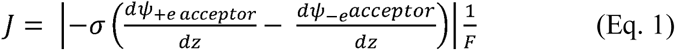

Where *dψ*+_*e-acceptor*_/*dz* is the linear gradient of the EP in the oxygen or nitrate zone measured in the presence of either of the two electron acceptors in the water column. *dψ*_*-e acceptor*_/*dz* is the gradient of EP in the same zone measured in the absence of electron acceptors. *F* is the Faraday’s constant, and σ is the sediment conductivity estimated from the conductivity of the sediment porewater and the porosity, according to (17). The conductivity of the porewater was measured with a conductivity meter (Mettler Toledo, Fisher Scientific, USA).

### ^15^N-based quantification of nitrate reduction pathways

The CB- and the CB-free cores in the ^15^NO_3_^−^-amended aquaria were equipped with top Plexiglas® tubes (Fig. 1C-E). The water enclosed in the tubes (app 150 ml) was kept mixed by a Teflon-coated magnetic stir bar suspended at a few centimeters above the sediment and driven by an external rotating magnet. The sediment cores with mounted tubes were maintained submerged inside the anoxic aquarium with the lids open and the stirrers on to assure initial homogenous conditions. Prior to the start of the incubation, water samples for analysis of nitrite, ammonium, and di-nitrogen isotopic composition were collected from the aquaria with a syringe. Samples for nitrite and ammonium analysis were transferred to plastic vials, immediately placed on ice and subsequently frozen (−20⁰C) until further analysis. Samples for di-nitrogen analysis were transferred into 12 ml Exetainers (Labco, U.K.) and fixed with 250 µL ZnCl_2_ (50:50 w/v). The tubes were then sealed with lids to initiate the incubation. After a 3 to 4-hour incubation period, the lids were removed and the water was sampled for nitrite, ammonium, and di-nitrogen determinations as described above. At the end of the sampling procedure, the cores were placed back in the anoxic aquarium for one to three days with the lid open before the incubation was repeated. The incubation procedure was repeated three times over the span of ten days to confirm stability of the system, thereby confirming that reaction rates can be approximated to the fluxes. Fluxes of N–isotopologues were calculated according to: *J*= (*C*_*end*_ - *C*_*start*_) / (*A x t*), where *C*_*end*_ and *C*_*start*_ are the concentrations of a given N-species in the overlying water at the end and at the start of the incubation, respectively; *A* is the surface area of the sediment core; and *t* is the incubation time. Rates of DNRA were calculated from the ^15^NH_4_^+^ flux divided by the fraction of [^15^NO_3_^−^] to total [NO_3_^−^] in the aquaria, *i.e.*, 0.45. Rates of denitrification were calculated from the fluxes of ^29^N_2_ and ^30^N_2_ according to (18). Rates of nitrite production were estimated from the fluxes of nitrite. The isotopic composition of di-nitrogen and ammonium was determined by mass spectrometry on a 20-22 hydra Isotope Ratio Mass Spectrometer (SerCon, Crewe, U.K.). [^29^N_2_] and [^30^N_2_] were determined as described in (19), while the ^15^N-atom% of ammonium was determined with the Combined Microdiffusion-Hypobromite Oxidation Method (20). [^15^NH_4_^+^] was calculated as the product of the ^15^N-atom% of ammonium and the ammonium concentration, the latter being determined with the Salicylate-Hypochlorite Method (21). Nitrite concentrations were determined by Ion Chromatography (Dionex IC-2500, Thermo Fisher Scientific).

### RNA sampling and extraction

Three CB-cores were sampled after the pre-incubation period (oxic controls) and three CB-cores were sampled after the anoxic ^15^NO_3_ ^−^ amended incubation. Sediment cores were extruded from the liners using a threaded rod. The upper 1 mm section from each core was transferred to 10 mL Falcon tubes and snap-frozen into liquid nitrogen. Total RNA from each sediment sample was extracted using the RNeasy PowerSoil Total RNA Kit (Qiagen). The RNA was concentrated with the kit RNA Clean & Concentrator (Zymo Research, USA) with an in-column DNase treatment (Ambion, USA) and stored at −80 °C before sequencing. Illumina sequencing (HiSeq, SR50 reads) was performed by DNASense (Aalborg, Denmark), after 16S and 23S rRNA removal with the Ribo-Zero rRNA Removal Kit for Bacteria (Illumina, USA).

### Transcriptomic analysis

The reads (approx. 10^7^ per sample) were mapped onto the genome of *Ca.* Electronema sp. GS (14) using the IMG annotation (taxon id 2728369268) and standard parameters in the software seal (https://jgi.doe.gov/data-and-tools/bbtools/bb-tools-user-guide/seal-guide/). The mapped reads were normalized to gene length and total read depth using the method reads per kilobase transcript per million mapped reads (RPKM) and ranked to determine the uniform expression levels. Differential expression between the nitrate incubations and the oxic controls was tested with the DESeq2 R package (22), with the false discovery rate (FDR) adjusted to 0.1. Analysis focused on genes presumably involved in nitrate and nitrite reduction (*napA* and gene neighbourhood) as proposed in (14). Transcriptomic data are available at NCBI/SRA under accession numbers PRJNA575156 and PRJNA575166.

### Phylogenetic analysis

Published *nrfA* (23) and *napA* (24) phylogenies were used to retrieve reference sequences for phylogenetic analyses of pMHC and *napA*, together with closest BLAST hits and, for pMHC, characterized, highly similar proteins from *Beggiatoa* (25) and Epsilonproteobacteria (εHao) (26). Reference sequences for *napD* and *napF* phylogenetic analysis were selected from the same genomes as *napA* when present. *napA*, *nrfA*/εHao/pMHC, *napD*, and *napF* were translated in silico, and amino acid sequences were aligned using the Muscle algorithm in Mega X version 10.0.5 with standard settings. Phylogenies were reconstructed using the maximum-likelihood algorithm in Mega X version 10.0.5 with a model of rate heterogeneity and the LG+G (NapA, NapD NapF) or WAG+G (pMHC) protein substitution model. The node stability was tested by bootstrapping.

## RESULTS

### The electric potential distribution in CB- and CB-free cores during pre-incubation

During the oxic pre-incubation, the EP at 1.5 cm depth increased progressively in the CB-cores to a maximum of 20 mV after 27 days (Fig. 2A). The magnitude of the electric field estimated from the gradient of the EP in the oxic zone (top 1 mm. Gradient calculated in the 0.2 to 1.5 mm depth interval) increased from 0.18 V m^−1^ to 1.8 V m^−1^ at week two and to 3.5 V m^−1^ at week four. This progressive increase in field strength confirmed that *Ca.* Electronema sp. GS was growing in the CB-cores. The EP was measured in the CB-free cores after two weeks of pre-incubation and reached 2.4 mV at 1.5 cm depth (Fig. 2B). The magnitude of the electric field in the 0.2-1.5 mm depth interval was 0.03 V m^−1^, corresponding to 2.5% of the field strength in the CB-cores. This low field strength was taken as evidence for insignificant presence of CB-in the CB-free cores, a pattern that was confirmed from microscope inspection prior to moving the cores into the anoxic, nitrate-amended aquaria.

**Figure 2.**
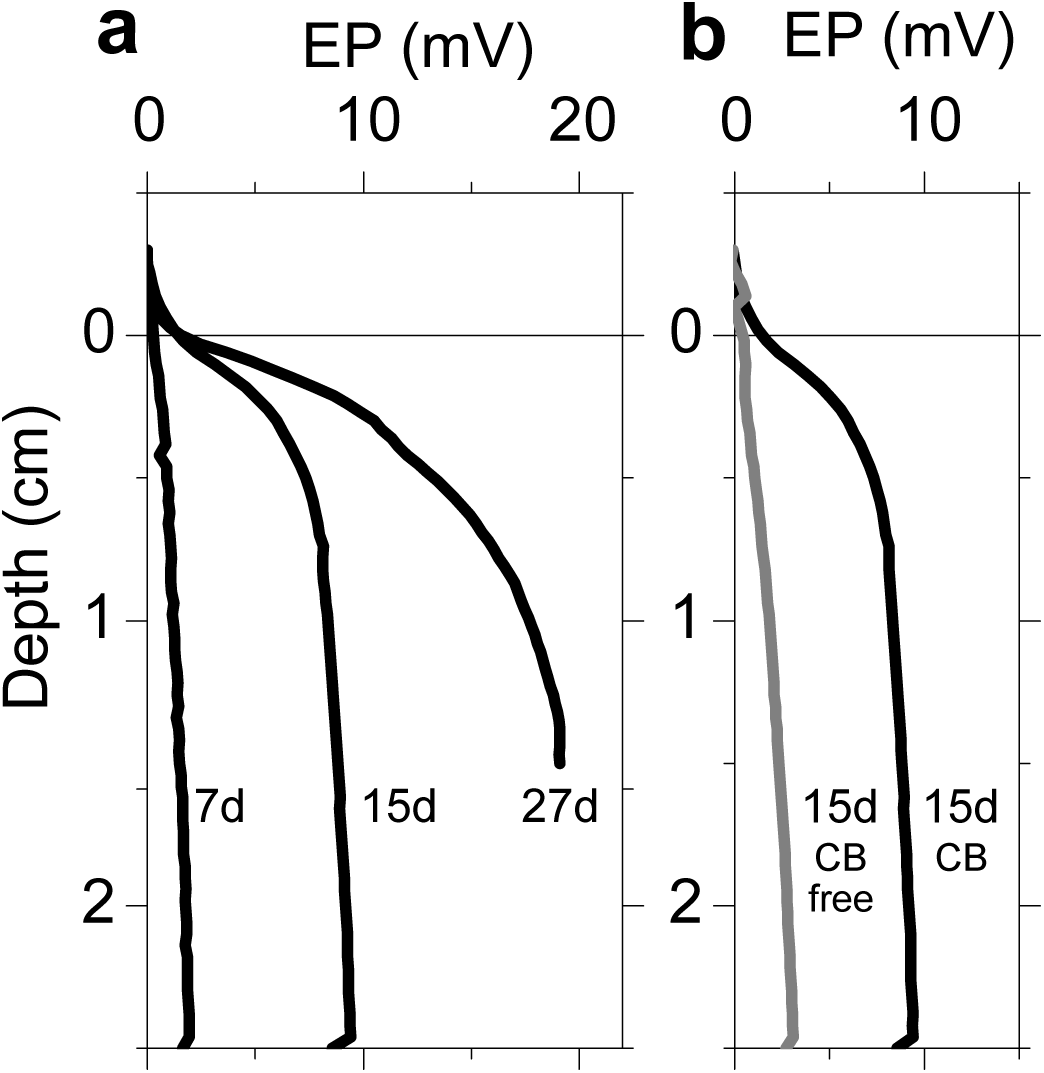
Electric potential (EP) depth profiles in sediment cores at day 7, 15, and 27 after the inoculation with cable bacteria (CB) (panel a). EP depth profiles in a sediment core at day 15 after the inoculation with cable bacteria (black line) and a CB-free core not inoculated (grey line) (panel b). Data are shown as mean (n = 3).

### Electron acceptor switch experiment with CB-cores

The EP distribution measured in CB-cores incubated for two days in anoxic, nitrate-amended water is shown in Figure 3. When the CB-core was exposed to air-saturated, nitrate-free water, the EP at 1 cm depth was 8 mV and the electric field in the 0-1.5 mm domain was 0.95 V m^−1^ or 27% of the field strength measured prior to the transfer, indicating a drop in CB activity. When the water above the sediment was turned anoxic, the EP at 1 cm depth and the electric field in the 0.2-1.5 mm domain was close to zero. When 200 µM nitrate was added to the anoxic water overlying the sediment, the EP at 1 cm depth and the electric field in the nitrate reduction zone (*i.e.*, 0 to <1.5 mm depth. Gradient calculated in the 1.2-2.4 mm depth interval) was similar to those measured in the presence of an oxygen saturated, nitrate-free water column, *i.e.*, 10 mV, and 1.0 V m^−1^ respectively. These observations suggest that the population of *Ca.* Electronema sp. GS was active both in the presence of an oxygen saturated, nitrate-free water column and in the presence of an anoxic nitrate-amended water column, but also that the activity of the population had declined significantly after being exposed to anoxia and nitrate for two days. The electron flux estimated from the EP profile measured in the presence of nitrate was 1.3 ± 0.1 mmol e^−^ m^−2^ h^−1^ (mean ± s.e.m., n = 3).

**Figure 3.**
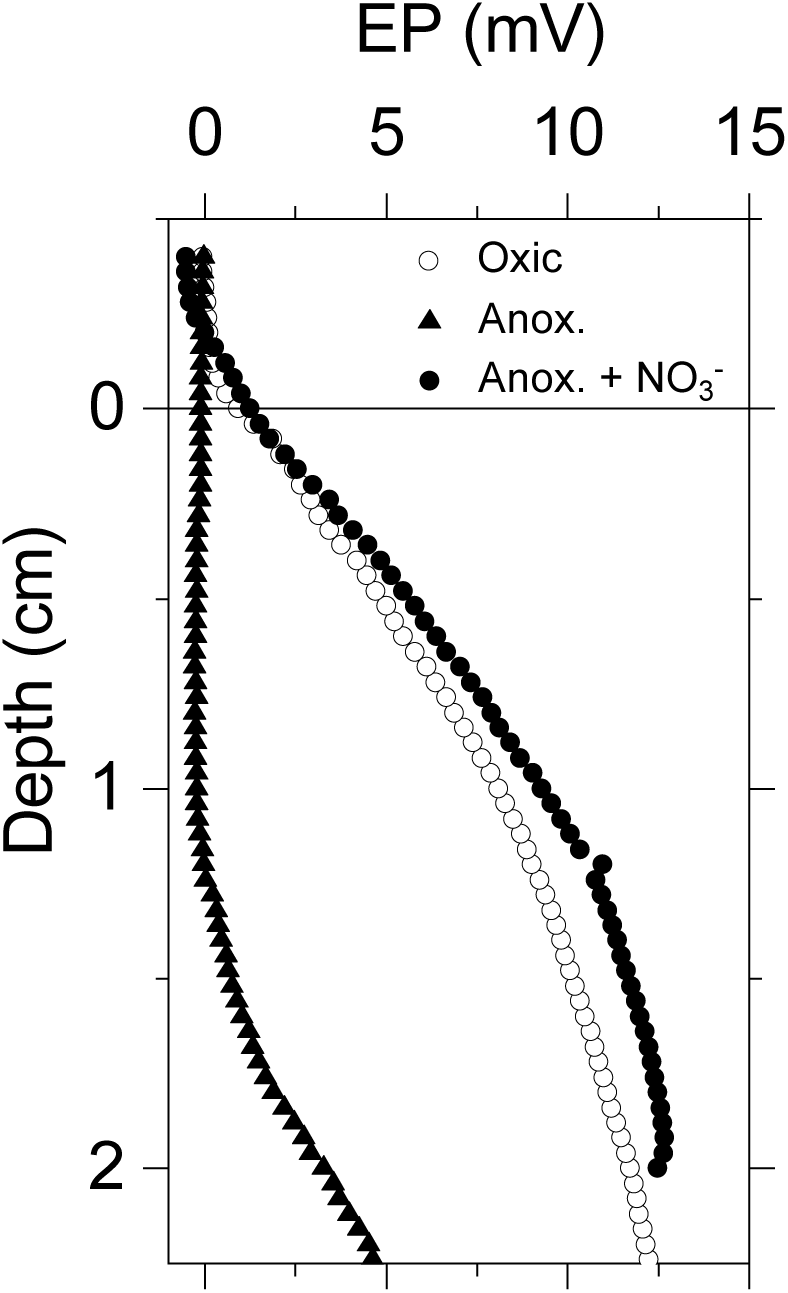
EP depth profiles in a CB-core sequentially exposed to aerated (white circles), anoxic, nitrate-free (black triangles), and anoxic nitrate-amended (black circles) water. Data are shown as mean (n = 3).

### ^15^N-based quantification of the nitrate reduction pathways

Rates of DNRA measured in the CB-cores were significantly different from those measured in the CB-free cores (t-test, p <0.01, n = 24) and up to two-fold higher (Figure 4). Rates of nitrite production in the CB-cores were significantly different from those measured in the CB-free cores (t-test, p <0.01, n = 24) and up to five-fold higher. There was no significant difference in denitrification rates between CB- and CB-free cores (t-test, p = 0.26, n = 24).

**Figure 4.**
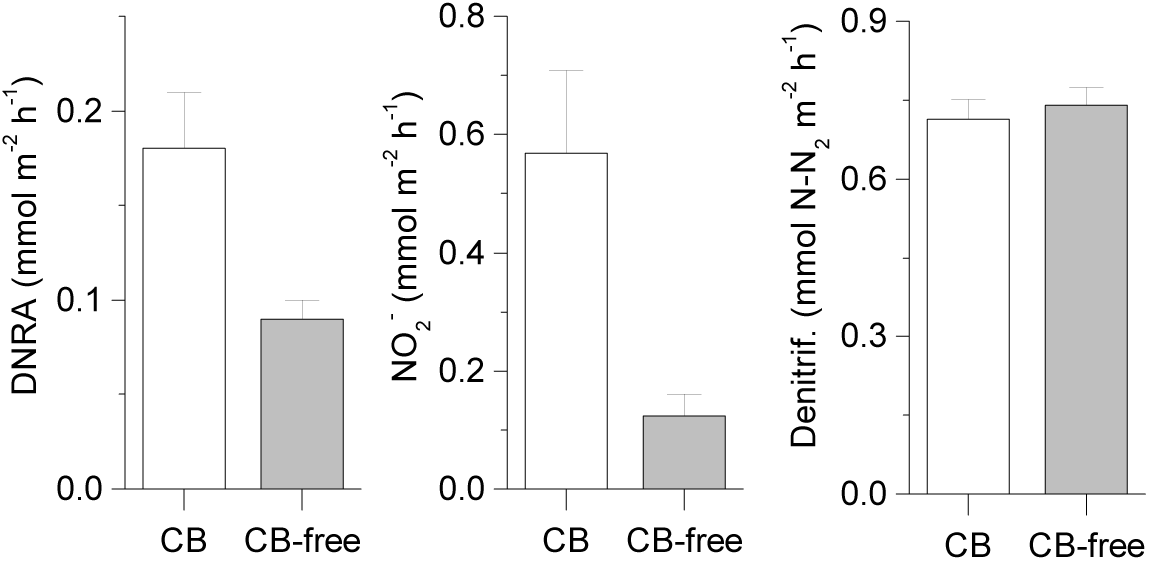
Net fluxes of NO_2_^−^, N_2_ from denitrification (D_w_) and NH_4_^+^ from DNRA across the sediment-water interface during anoxic incubations of sediment inoculated with cable bacteria and in control cores. Data are shown as mean ± s.e.m. (n = 12).

### Operon structure, expression, and phylogeny of putative DNRA genes

Detailed analysis of the gene neighborhood of the *napAB* genes revealed that *Ca.* Electronema sp. GS contained a complete *nap* operon encoding all components for periplasmic reduction of nitrate to nitrite (Fig 5A): the maturation factor NapF, two copies of the chaperone NapD, the electron-transferring membrane complex NapGH, and the catalytic complex NapAB. The gene encoding a pMHC, hypothesized to reduce nitrite to ammonium, is located directly downstream of the *nap* operon (Fig. 5A). All eight genes were among the 100 most highly expressed genes (out of 2,649 total genes) under nitrate-reducing conditions (Fig. 5A; Table S1). A comparison of gene expression levels between the nitrate-amended incubations and the oxic controls showed that all eight consecutive genes were slightly but consistently upregulated under nitrate-reducing conditions (Fig. 5B). However, according to DESeq2 analysis, none of these upregulations were significant (Table S1).

**Figure 5.**
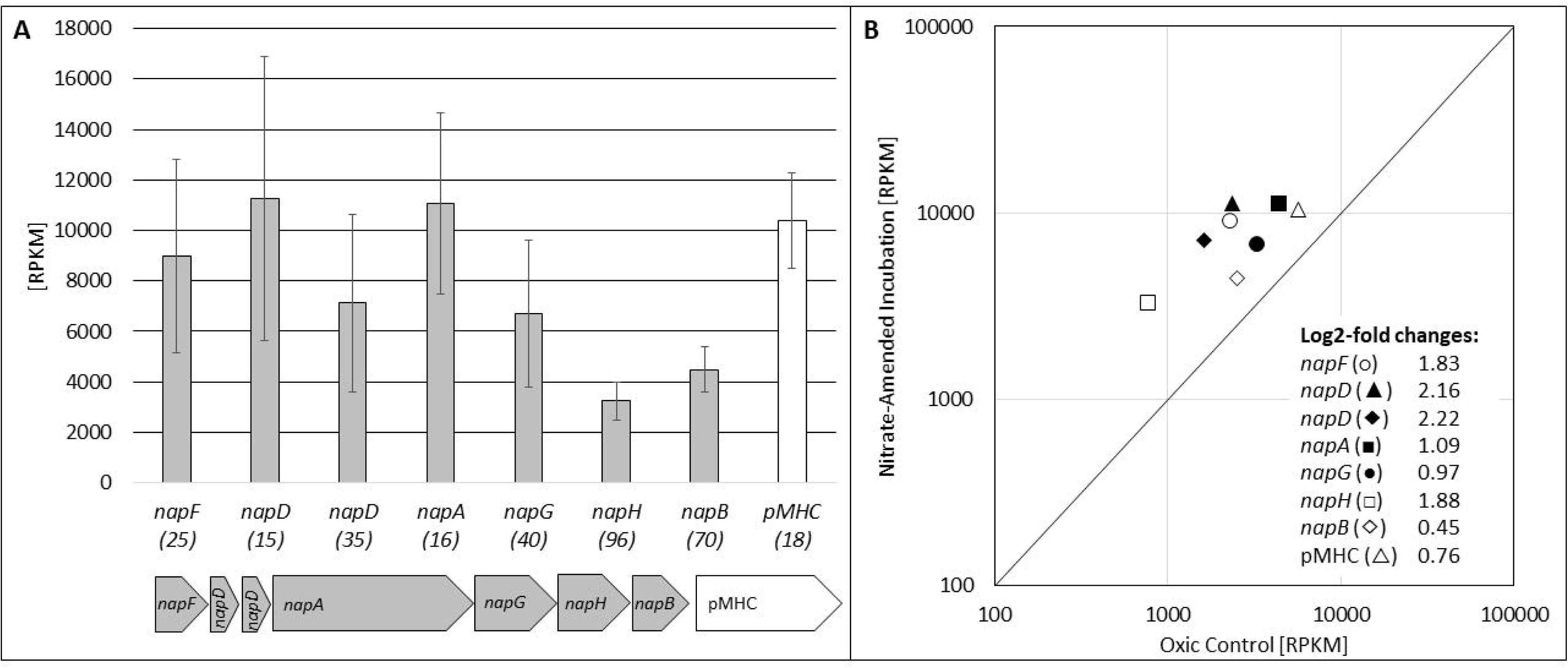
Structure and gene expression of the putative DNRA genes in *Ca.* Electronema sp. GS. **(A)** Operon structure and expression levels under nitrate-reducing conditions (n=3, ± s.e.m.). The rank of each gene in the transcriptome is shown in parenthesis. RPKM: <u>R</u>eads <u>P</u>er <u>K</u>ilobase per <u>M</u>illion. **(B)** Expression of the *nap*-pMHC operon after nitrate-amendment compared to the oxic controls (n=3).

BLAST analyses retrieved closest matches for *napF* (44.9% translated amino acid identity), *napD* (50.6 and 43.2%), and pMHC (73.8%) to homologs in the marine cable bacterium *Ca.* Electrothrix aarhusiensis MCF (14), and additional matches for pMHC (70.0%) and *napF* (53.1%) to gene fragments in the less complete genomes of *Ca*. Electrothrix marina A3 and A5, respectively. No other DNRA genes were at first detected in cable bacteria but a closer inspection of published cable bacteria genomes revealed that *napF* in *Ca.* E. aarhusiensis MCF is located at the end of a contig, and a short (320 amino acids) *napA* fragment is located at the end of another (short) contig in *Ca.* E. aarhusiensis MCF, which together suggests that the rest of the *nap* operon was missed during sequencing/genome assembly in this marine cable bacterium. Phylogenetic analysis placed the cable bacterial NapA, NapD, and NapF lineages in clades containing different alpha-, beta-, or gammaproteobacterial genes (Figures S1, S2, S3). The cable bacterial pMHC formed a monophyletic clade with homologs from sediment-dwelling bacteria of at least three phyla (Nitrospirae, Calditrichaeota, and Gammaproteobacetria, which include large sulfur bacteria; Fig. 6 and Fig. S4). This pMHC clade furthermore forms a sister clade to proteins of the epsilonproteobacterial hydroxylamine oxidoreductase (εHao) family (26).

**Figure 6.**
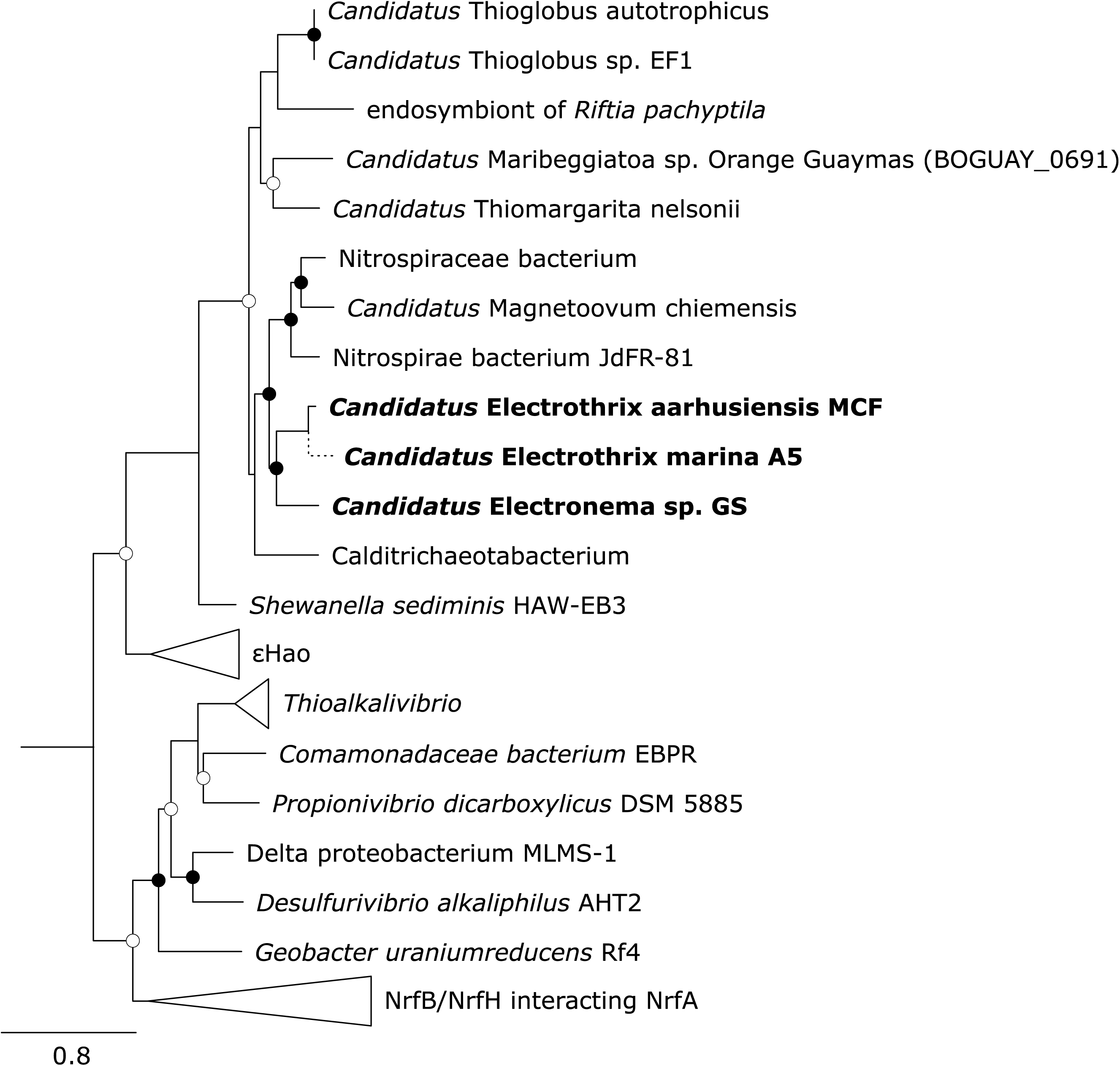
Phylogenetic affiliation of cable bacterial periplasmic multiheme cytochromes (pMHC) with εHao and NrfA proteins. Consensus tree build by maximum likelihood and supported with 1000 bootstrap resamplings. Bootstrap values are represented by circles: open >70%, filled >90%. Tree was rooted with the nitrite reductase (nirB) of *Bacillus subtilis*. Scale bar represents 0.8 estimated amino acid substitutions. The phylogenetic position of the short NapF fragment of *Ca*. E. marina was calculated by the maximum parsimony method in the program ARB (43) without changing the overall tree topology. For an expanded phylogenetic tree, see Figure S4.

## DISCUSSION

### *Ca.* Electronema sp. GS employs the DNRA pathway

Results from the ^15^N experiment showed similar rates of denitrification in CB- and CB-free cores suggesting that *Ca.* Electronema sp. GS does not reduce nitrate to N_2_, which is in line with previous studies showing lack of EP responses to nitrous oxide amendments in sediments with marine cable bacteria (12). The higher rates of DNRA and anaerobic nitrite production in CB-cores compared to CB-free cores suggest that *Ca*. Electrononema sp GS. can reduce nitrate to nitrite and reduce nitrite further to ammonium. This capacity for nitrate and nitrite reduction aligns with previous studies of marine cable bacteria (11, 12), showing that these can reduce nitrite and possibly nitrate, though the end product of nitrite reduction was not identified in these studies.

Assuming that the difference in rates of DNRA and nitrite production between CB- and CB-free cores reflects the nitrate reduction activity of the *Ca.* Electronema sp. GS population, 446 ± 145 µmol NO_2_^−^ was produced per m^2^ per hour from nitrate reduction, and 89.7 ± 33 µmol NH_4_^+^ was produced per m^2^ per hour from DNRA. The electron flux from electron donors needed to sustain these rates can be estimated from the stoichiometry of the following cathodic reactions:

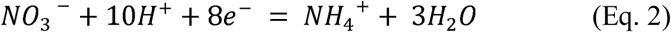

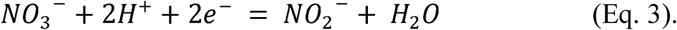

The so calculated electron demand amounted to 1.6 ± 0.4 mmol e^−^ m^−2^ h^−1^. The flux of electrons supplied in the nitrate reduction zone by the deeper electrogenic oxidation of sulphide, estimated from the EP profiles in Figure 3, amounted to 1.3 ± 0.1 mmol e^−^ m^−2^ h^−1^. The close match between these independent estimates of electron demand and supply further strengthen our hypothesis that the freshwater cable bacterium *Ca.* Electronema sp GS can reduce nitrate to nitrite and nitrite further to ammonium, via e-SOx.

This hypothesis is further supported by our genomic and transcriptomic data. Periplasmic reduction of nitrate to nitrite can be deduced from the presence and expression of a complete *nap* operon in *Ca.* Electronema sp. GS (Fig. 5), where the only unusual feature appears to be a duplication of the chaperone-encoding *napD* (Fig. S2; (27)). Identification of the nitrite reductase has been more challenging but several lines of evidence point at the pMHC for nitrite reduction to ammonium: (i) the pMHC is closely related (57% amino acid identity; Fig. 6) to a multiheme cytochrome (BOGUAY_0691) of orange *Ca.* Maribeggiatoa sp., which has been functionally characterized and can reduce nitrite to ammonium (15, 25); also the εHao sister clade to the pMHC (Fig. 6) has been experimentally proven to reduce nitrite to ammonium (26); (ii) the pMHC is positioned downstream of the *nap* operon, and the entire *nap*-pMHC gene structure is syntenic to that of the well-characterized DNRA genes of epsilonproteobacteria (26), suggesting a linked function for DNRA; (iii) the pMHC is, together with the entire *nap* operon, highly expressed under nitrate-reducing conditions (Fig. 5A, Table S1); and (iv) although not significant for any single gene, all 7 *nap* genes and the pMHC are slightly but consistently upregulated under nitrate-reducing compared to oxic conditions (Fig. 5B, Table S1). Taken together, our data support a model in which nitrate is first reduced to nitrite by NapAB, and then further reduced to ammonium by pMHC.

### Origin and distribution of DNRA in cable bacteria

Cable bacteria, including *Ca.* Electronema sp. GS, are a monophyletic sister group to the genus *Desulfobulbus* within the deltaproteobacterial Desulfobulbaceae. In contrast, none of their *nap* and pMHC genes were phylogenetically affiliated with deltaproteobacterial genes (Fig. 6; Figures S1-S4). This indicates that all genes for DNRA have been acquired by horizontal gene transfer, and since the various genes show distinct evolutionary histories, probably in multiple events from different organisms. Possible donors for the different *nap* genes are different members of the Alpha-, Beta-, and Gammaproteobacteria (Figures S1-3). For the pMHC, it is intriguing that the most closely related genes were from bacteria that can co-occur with cable bacteria in sulphide-oxygen gradients of surface sediments, like the large sulphur bacteria *Thiomargarita* or *Beggiatoa* (9, 28), Calditrichaeota (29), or the magnetotactic bacterium *Magnetoovum* (30). This implies that horizontal gene transfer may occur between such phylogenetically distant organisms in this shared niche.

We can currently not infer how widespread DNRA is among cable bacteria. The only complete DNRA pathway was detected in our model freshwater species *Ca.* Electronema sp. GS (Fig 5A), while DNRA genes were rare and fragmented in the marine genus *Ca.* Electrothrix. Whether this is due to its incomplete and highly fragmented genomes (see results above and (14)) or due to its truly limited potential for DNRA cannot be resolved. It is however striking that the few DNRA genes detected in *Ca.* Electrothrix (*napF*, *napD*, pMHC) always cluster with their freshwater homolog in *Ca.* Electronema sp. GS (Fig. 6, Figures S1-4). As crossovers between freshwater and marine species are rare (31), this indicates that the capacity for DNRA was acquired by cable bacteria before their diversification into different salinity niches represented by the candidate genera Electrothrix and Electronema (32). This scenario suggests that all cable bacteria originally were capable of DNRA, but the pathway (or part of it) may subsequently have been lost in certain species or lineages, as is commonly observed for canonical nitrate reducers and denitrifiers (33–35). The observation of nitrate-dependant e-SOx in marine systems (11, 12) supports an extant capability of DNRA across the genus divide.

### The architecture of the nitrate reduction system of *Ca*. Electronema

On the basis of the transcriptome data and the morphology of cable bacteria a hypothetical model of the architecture of the nitrate reduction system in *Ca*. Electronema sp. GS can be described (Fig. 7). *Ca*. Electronema sp. GS expressed a complete *napFDDAGHB* operon, as well as genes encoding a putatively nitrite-reducing pMHC. The general arrangement and biochemical role of the proteins produced from the *napFDDAGHB* transcripts can be drafted from the model of the respiratory Nap system of the epsilonprotobacterium *Wolinella succinogenes* (27, 36). According to this model, the nitrate-reducing NapAB complex is located in the periplasm. The NapGH complex is embedded in the inner membrane and catalyses the oxidation of reduced menaquinonens dissolved in the inner membrane, passing the electrons to NapA via NapB. According to Simon and Klotz (27), the NapGH complex transfers the protons liberated from menaquinone oxidation to the periplasm. The NapD is a chaperone protein that assists the transfer of mature NapA via the TAT complex (also present and expressed in *Ca*. Electronema sp. GS, data not shown) from the cytoplasm to the periplasm, whereas the NapF protein, located in the cytoplasm is proposed to provide reducing power for the maturation of NapA (27). In *W. succinogenes*, nitrate reduction via the Nap system involves the reduction of menaquinones by a membrane-bound formate dehydrogenase (FdhABC) (36), and a proton motive force (*pmf*) is supposedly generated through the NapGH-FdhABC mediated redox cycle of the menaquinones (36). If the Nap system of *Ca*. Electronema sp. GS should work in a similar way, but driven by e-SOx, electrons for menaquinone reduction should be delivered from the conducting strings present in the periplasm (2) via, *e.g.*, the highly expressed periplasmic cytochromes (14, 37) and a hitherto unidentified membrane-bound menaquinone reductase (like, *e.g.*, the Type I cytochrome c3:menaquinone oxidoreductase protein Qrc found in *Desulfovibrio vulgaris* Hildenborough (38)). E-SOx-mediated nitrate reduction via the conventional Nap pathway would further imply electron transport from anodic cells to cathodic cells both having electron carrier molecules with identical standard redox potential, as menaquinones are also implicated in cytoplasmic redox cycling during anodic sulphide oxidation in *Ca*. Electronema sp. GS (14). Modelling such transport from a concentration cell analog (SI Extended discussion) shows that such an electron flow is thermodynamically feasible, and that the voltage produced can be sufficiently high to drive an electric current that exceeds the current reported for cable bacteria.

**Figure 7.**
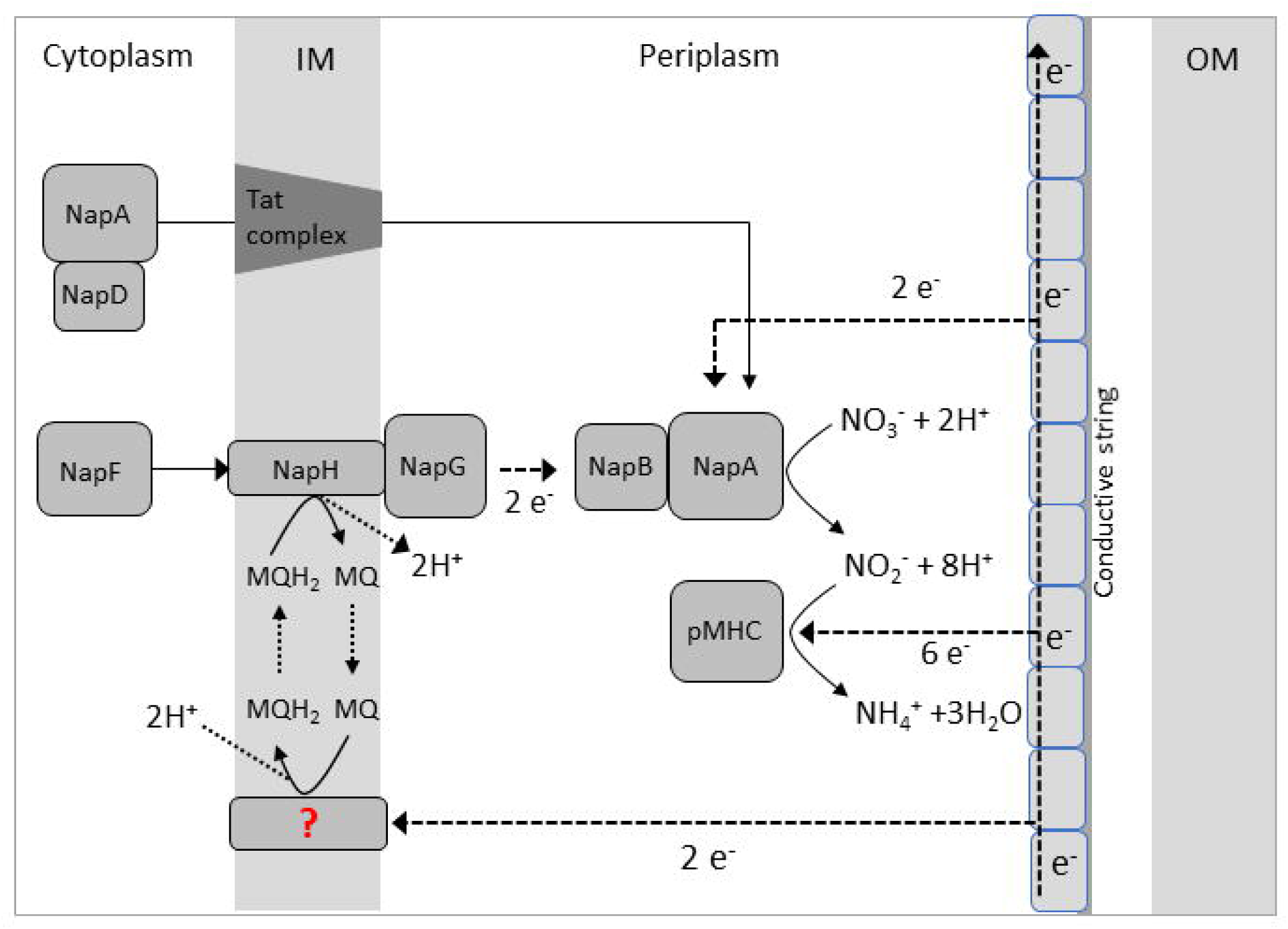
Model of cathodic nitrate reduction to ammonium in *Ca.* Electronema sp. GS. Dotted arrows indicate electron flow. Electrons are derived from anodic sulfide oxidation and delivered via the periplasmic conductive fiber to the periplasmic NapAB complex and pMHC. Electron flow could occur merely in the periplasm or involving menaquinone cycling and the membrane-bound NapHG complex.

The putative nitrite reductase (pMHC) of *Ca*. Electronema sp. GS has presumably a periplasmic localization, as also suggested for its close homolog in *Ca.* Maribeggiatoa sp. (15). The pMHC-based nitrite reduction system in *Ca.* Maribeggiatoa sp is not described. From the genome data, we cannot identify an electron donor that links nitrite reduction to menaquinone cycling, as homologs to, *e.g.*, the NrfHA system in Epsilonproteobacteria (39), the NrfBCD system in Gammaproteobacteria (27), and the CymA system in *Shewanella oneidensis* (40, 41) are absent in CB. It is possible that nitrite reduction in *Ca*. Electronema sp. GS is fully periplasmatic and decoupled from menaquinone cycling. In this scenario the pMHC would receive electrons directly from the conducting strings or via the periplasmic cytochromes.

In *W. succinogenes*, proton production from FdhABC-mediated formate oxidation matches the proton consumption from periplasmatic nitrate reduction (36). NapGH-mediated proton transfer into the periplasm could therefore in principle produce a *pmf*, which could drive ATP synthesis through oxidative phosphorylation. The generation of a *pmf* by NapGH in *Ca*. Electronema sp. GS is however less likely, as protons from electron donor oxidation are produced by distant anodic cells in the filament (1, 3), implying proton deficiency in the periplasm of the nitrate-reducing cathodic cells and consequently loss of *pmf* through proton consumption by periplasmic nitrate reduction. Alternative electron flows, such as direct electron transfer from the conductive strings via periplasmic cytochromes to the NapAB complex or to pMHC would not generate a *pmf* during nitrate/nitrite reduction either, as these enzyme-mediated reactions take place in the periplasm, disconnected from the cytoplasmic membrane. In conclusion, cathodic cells performing nitrate reduction cannot conserve energy through oxidative phosphorylation. This matches the situation proposed for oxygen-reducing cathodic CB cells, which appear to perform periplasmic oxygen reduction without energy conservation (14).

## Conclusions and perspectives

On the basis of geo(electro)chemical and transcriptomic evidences we conclude that *Ca*. Electronema sp. GS reduces nitrate via the DNRA pathway. The reduction of nitrate to nitrite and nitrite to ammonium is catalyzed by a NapAB complex and a pMHC, respectively. The capability for DNRA has likely been acquired by horizontal gene transfer from phylogenetically distant organisms that share the same ecological niche. The periplasmic allocation of these reductases implies no energy conservation via oxidative phosphorylation in association with nitrate/nitrite reduction, supporting the hypothesis of a division of labor along the filament, with the cathodic cells serving primarily as “flare” for electrons delivered by the anodic cells. In this scenario, high energy-demanding transcription and protein synthesis occur primarily in the suboxic zone (14), and the movement of CB filaments (42) across the sediment geochemical gradient alternates the exposure of CB cells to sulphide-oxidizing and nitrate-reducing conditions; the limited ATP demand for protein repair (e.g. by chaperones) and maintenance in the cathodic cells may be provided from storage compounds (polyphosphate, polyglucose (14)) produced under anodic conditions. The apparent duplication of the chaperon napD (Fig. S2) may be an adaptation to this lifestyle.

From a geochemical perspective, our results indicate that sediment colonization by CB would favor the recycling of bioavailable nitrogen via DNRA over its release to the atmosphere via denitrification. Cable bacteria have also been reported to stimulate ammonium production in sediment overlaid by oxic water, via promoting acid dissolution of FeS minerals and subsequent DNRA activity by Fe^2+^ (or H_2_S) oxidizing chemolithotrophs (13). Assuming that CB will favor O_2_ over nitrate as terminal electron acceptor, the predominance of the direct or indirect stimulation of DNRA by CB will ultimately depend on the availability of O_2_, Fe^2+^ (dissolved or precipitated in FeS minerals), and organic carbon (OC), with the ratio between Fe^2+^ and OC determining the relative importance of heterotrophy over Fe-fuelled chemolithotrophy. Indirect stimulation of DNRA by CB should thus dominate in oxic sediments with relatively high Fe:OC ratios, whereas the direct effect would prevail under O_2_-depleted, nitrate- and organic-rich conditions. Suitable sites for the latter are sediments of eutrophic water bodies with transiently or permanently anoxic bottom water. As both the direct and indirect effects of e-SOx converge in stimulating DNRA, CB can potentially play an important and overlooked role in regulating the balance between denitrification and DNRA in sedimentary environments. In particular, the contribution of CB to N transformation might have gone unaccounted in sediment slurry incubations, where the chemical gradients that favor e-SOx (*i.e.*, physical separation between electron acceptors and donors) are artificially broken to maximize mass transport.

## ACKNOWLEDGMENTS

We are grateful to Lars B. Pedersen for microsensor construction, to Jeanette Johansen, Susanne Nielsen, and Karina B. Oest for help with laboratory analysis. Tage Dalsgaard is acknowledged for support to the isotopic analysis, and Filip Meysman for fruitful and inspirational discussion. Results incorporated in this study have received funding from the European Union’s Horizon 2020 research and innovation programme under the Marie Sklodowska-Curie grant agreement No 656385 to UM; and the Danish National Research Foundation (grant DNRF104; NR-P, AS).

## COMPETING INTERESTS

Authors declare no competing interests in relation to the work described.

