## Supplementary material for "Dissimilatory Nitrate Reduction to Ammonium in the Cable Bacterium *Ca*. Electronema Sp. GS": SI

### Supplementary information I

Ugo Marzocchi[1,2,3*], Casper Thorup[2,3], Ann-Sofie Dam[2,3], Andreas Schramm[2,3], and Nils Risgaard-Petersen[2,3]

[1] Department of Chemistry, Vrije Universiteit Brussel, Belgium

[2] Center for Geomicrobiology, Section for Microbiology, Department of Bioscience, Aarhus University, Aarhus, Denmark

[3] Center for Electromicrobiology, Section for Microbiology, Department of Bioscience, Aarhus University, Aarhus, Denmark

Index:

**S1.** NapA phylogeny

**S2**. NapD phylogeny

**S3**. NapF phylogeny

**S4**. pMHC extended phylogeny

**S5**. Extended discussion

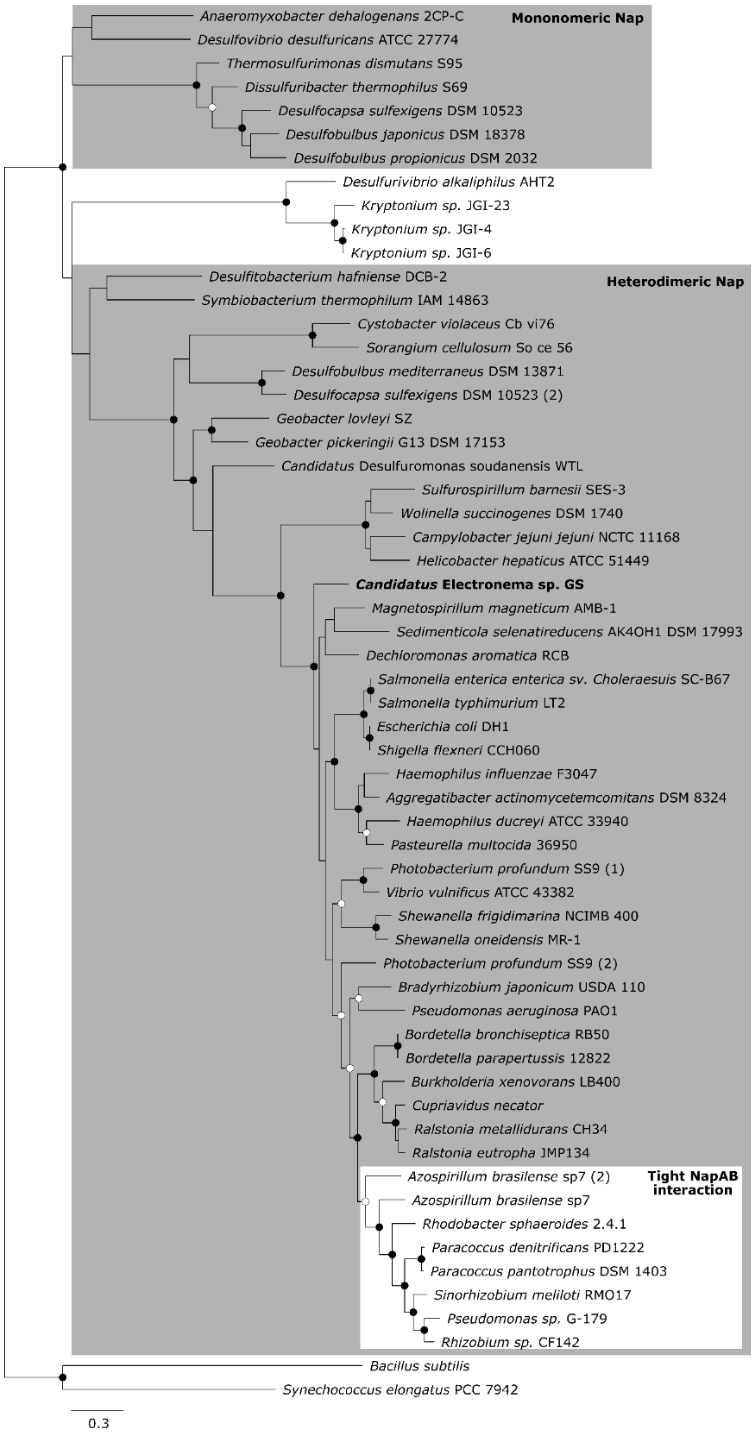


**Figure S1**. Phylogeny of the *napA* gene of *Ca.* Electronema sp. GS. Maximum likelihood tree supported by 1000x bootstrap resampling. Bootstrap values are represented by circles: open >70%, filled >90%. The scale bar represents 0.3 estimated amino acid substitutions. Tree was rooted with the nitrate reductase (nas) of *Bacillus subtilis*.

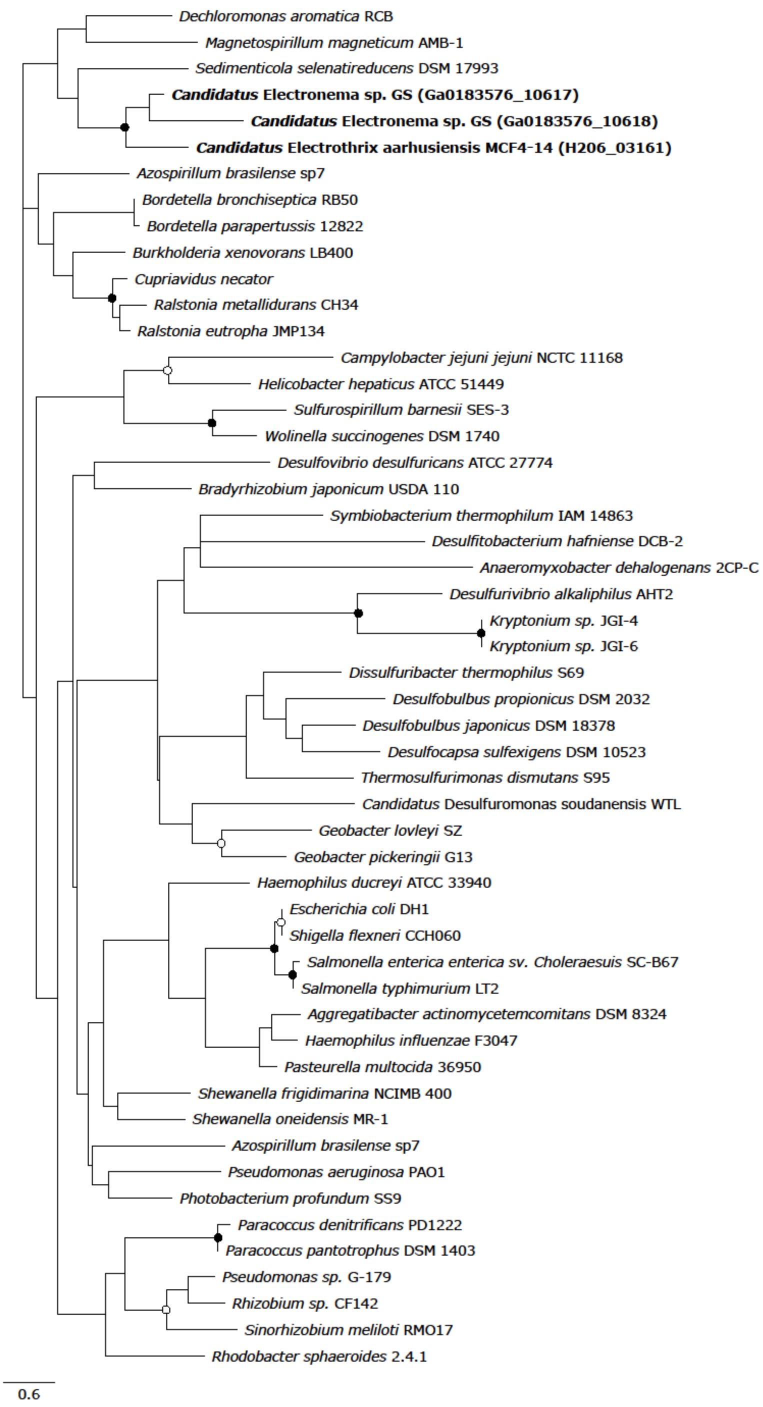


**Figure S2**. Phylogeny of the *napD* gene of *Ca*. Electronema sp. GS and *Ca*. Electrothrix aarhusiensis MCF. Maximum likelihood tree supported by 1000x bootstrap resampling. Bootstrap values are represented by circles: open >70%, filled >90%. The scale bar represents 0.6 estimated amino acid substitutions. Locus tags of cable bacteria genes are in parentheses.

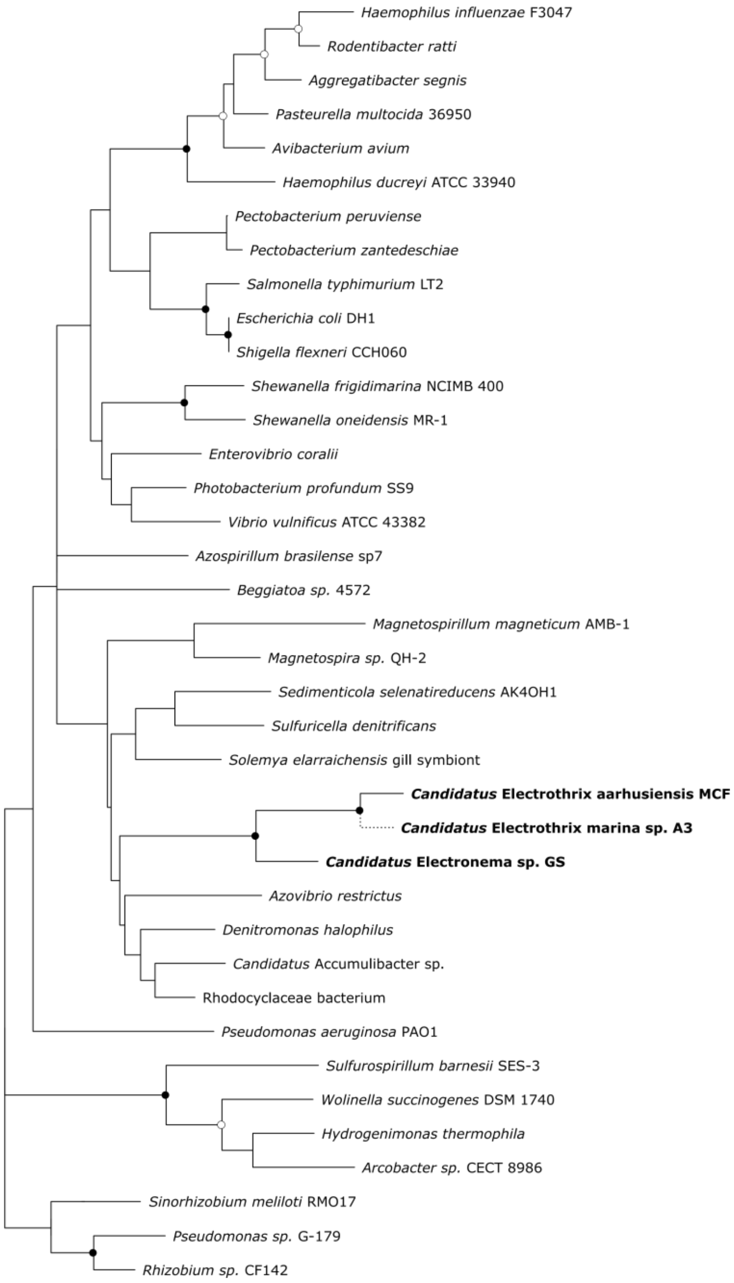


**Figure S3.** Phylogeny of the *napF* gene of *Ca.* Electronema sp. GS, *Ca.* Electrothrix aarhusiensis MCF and *Ca.* Electrothrix marina A5. Maximum likelihood tree supported by 1000x bootstrap resampling. Bootstrap values are represented by circles: open >70%, filled >90%. The scale bar represents 0.3 estimated amino acid substitutions. The phylogenetic position of the short NapF fragment of *Ca*. E. marina (dotted line) was calculated by the maximum parsimony method in the program ARB without changing the overall tree topology.

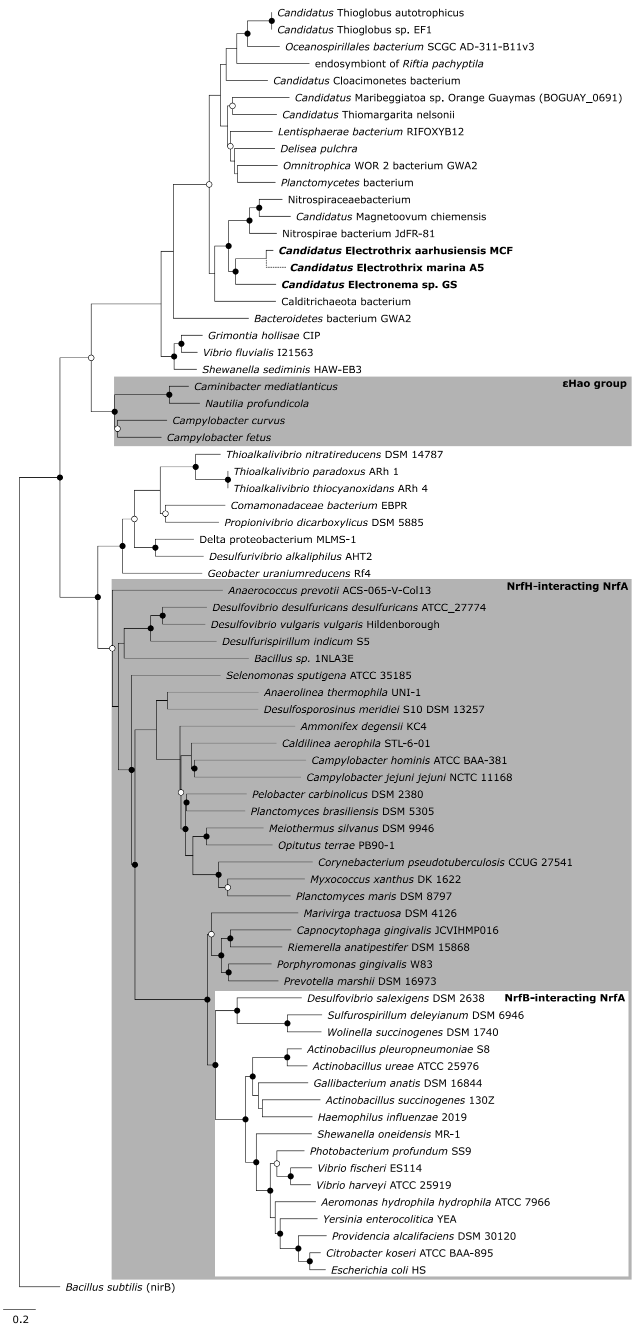


**Figure S4**. Full phylogeny of the pMHC of *Ca*. Electronema sp. GS and *Ca*. Electrothrix aarhusiensis MCF. Maximum likelihood tree supported by 1000x bootstrap resampling. Bootstrap values are represented by circles: open >70%, filled >90%. The scale bar represents 0.2 estimated amino acid substitutions. The phylogenetic position of the short pMHC fragment of *Ca*. E. marina (dotted line) was calculated by the maximum parsimony method in the program ARB without changing the overall tree topology. Tree was rooted with the nitrite reductase (nirB) of *Bacillus subtilis*.

S5. Extended discussion

### LINKING SULFIDE OXIDATION TO NITRATE REDUCTION VIA MENAQUINONE CYCLING THROUGH LONG-DISTANCE ELECTRON TRANSFER IN CABLE BACTERIA

The metabolic model of nitrate reduction in the cathodic nitrate-reducing cells in cable bacteria includes a menaquinone cycle in which a reduced menaquinone donates two electrons to the menaquinol dehydrogenase NapGH, membrane component NapH. This results in the transfer of two protons from the cytoplasm to the periplasm. The menaquinone is reduced back via sulfide oxidation, which however takes place in a distant cathodic cell, most likely via reverse sulfate reduction (1). Assuming the presence of an identical membrane-bound enzymatic apparatus for catalyzing menaquinone oxidation-reduction reactions in the cathodic nitrate-reducing cell and in the anodic sulfide-oxidizing cell, and that these apparatus are electrically connected with a conductor the metabolic system linking sulfide oxidation to nitrate reduction via a menaquinone cycle can be considere as a concentration cell (Fig. S5.1). This is a galvanic cell that has two equivalent half-cells with the same reactants differing only in concentrations. Such a concentration cell produces a voltage as it attempts to reach chemical equilibrium, which occurs when the concentration of the reactant in both half-cells are equal. This is achieved by transferring electrons from the half-cell with the highest concentration of reduced compounds to the half-cell with a lower concentration of these. In the following we will investigate, the thermodynamic constrains on the concentration cell model of nitrate reduction in cable bacteria in order to elucidate if the flow of electrons between the menaquinone in pool anodic (half) cells and the menaquinone pool in cathodic (half) cells via the conducting fibre in cable bacteria is thermodynamic possible and can account for current observed running in these organisms.

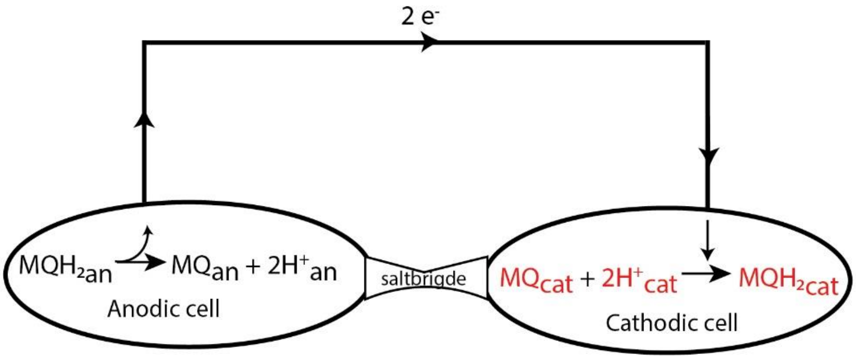


***Figure S5.1***. *Conceptual scheme of a concentration cell.*

**The voltage yield of the concentration cell**

The voltage produced by the concentration cell can be estimated from the Nernst equation:

$$E = E_o - \frac{0.0592}{n} LogQ \qquad \text{(Eq. S1)}$$

Where $E_0$ is the standard cell potential, which is defined as the sum of the reduction potential and the oxidation potential of the half reactions. $n$ is the number of moles of electrons transferred, and $Q$ is the reaction coefficient. The reduction potential ($E_{red}$) for the reaction MQ + $2H^+$ +$2e^-$ → $MQH_2$ is -67 ± 10 mV (2) and the oxidation potential is consequently +67 ± 10 mV. Hence, $E_o$ equals zero. The full reduction of 1 mole of menanquinones implies the transfer of two moles of electrons and $n$ consequently equals 2. The reaction coefficient $Q$ is defined from the overall reaction:

$$MQH_{2\,an} + MQ_{cat} + 2H^+_{cat} \rightarrow MQ_{an} + 2H^+_{an} + MQH_{2cat}$$

Thus:

$$Q = \frac{[MQH_{2cat}][MQ_{an}][H^+{}_{an}]^2}{[MQH_{2an}][MQ_{cat}][H^+{}_{cat}]^2} \qquad \text{(Eq. S2)}$$

The proton source for the menaquinone cycle is the cytoplasm. As an equivalent pH of the cytoplasm of cathodic and anodic half-cells can be assumed, $[H^+{}_{an}]^2/[H^+{}_{cat}]^2$ approximates 1. Hence:

$$Q \approx \frac{[MQH_{2cat}][MQ_{an}]}{[MQH_{2an}][MQ_{cat}]} \qquad \text{(Eq. S3)}$$

With *a* and *b* denoting the fraction of oxidized menaquinones (MQ) to the total pool of menaquinones ($MQ_{Tot}$) in the anodic cell and cathodic cell, respectively we have:

$MQ_{an}$ = a $MQ_{Total\ anode}$ (Eq. S4)

$MQH_{2an}$ = (1-a) $MQ_{Total\ anode}$ (Eq. S5)

for the anodic cell, and:

$MQ_{cat}$= b $MQ_{Total\ cathode}$ (Eq. S5)

$MQH_{2cat}$ = (1-b) $MQ_{Total\ cathode}$ (Eq. S7)

for the cathodic cell. Thus:

$$Q = \left( \frac{[(1-b)\ MQTotal_{cathode}\ ][a\ MQTotal_{anode}]}{[(1-a)\ MQTotal_{anode}\ ][b\ MQTotal_{cathode}\ ]} \right) = \left( \frac{(1-b)\ a}{(1-a)\ b} \right) \qquad \text{(Eq. S8)}$$

With this expression for *Q* the voltage produced by the concentration cell is given by:

$$E = -\frac{0.0592}{2} Log \left( \frac{(1-b)\ a}{(1-a)\ b} \right) \qquad \text{(Eq. S9)}$$

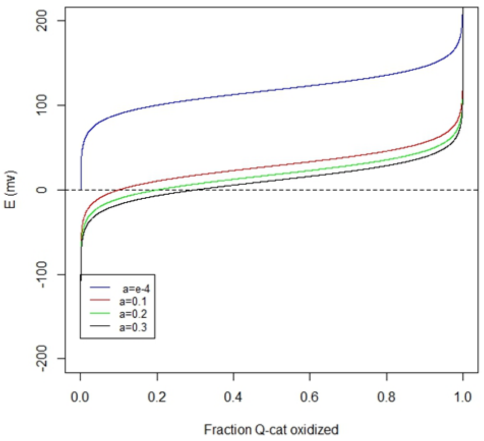

***Figure S5.2****. The voltage yield (E) of the concentration cell for different values of a (the fraction of oxidized menaquinonens in the anodic half-cell) as function of the fraction of menaquinons oxidized in the cathodic half-cell (b).*

Equation S9 allows to calculate the voltage yield ($E$) of the concentration cell for different values of $a$ and $b$ A positive voltage yield ($E > 0$) implies that the concentration cell operates in the forward direction (electron transfer from the anodic half-cell to cathodic half-cell), whereas a negative voltage yield implies electron transfer in the opposite direction. As shown in Figure S5.2, electron transfer from the menaquinone pool in an anodic half-cell to the menaquinone pool in a cathodic half-cell is most favorable if the electrons are delivered from an anodic half-cell with an almost completely reduced (99.99%, a = 0.0001) menaquinone pool. For such a system, electron transfer in the forward direction is possible if only 0.01% (b = 0.0001) or more of the menaqinone pool is kept oxidized in the cathodic half-cell. The thermodynamic threshold increases when electrons are delivered from anodic half-cells with a less reduced menaquinone pool. If electrons are delivered from anodic half-cells having 90% of the menaquinone pool reduced the recipient half-cell should keep more than 10% of the menaquione pool oxidized for the reaction to proceed, and for anodic cell having 70% of their menaquinone pool reduced, the recipient cell should keep more than 30% of its menaquinone pool oxidized, etc. The voltage yield is highest if electrons can be delivered from an anodic half-cell having an almost fully reduced menaquinone pool, and for such a system the voltage produced falls in the range 80-150 mV for 5% < b < 92%.  The voltage yield drops significantly when the menaquinone pool in the anodic half-cell becomes more oxidized. Figure S5.3 shows the voltage yield for a concentration cell, where 90% of the menaquinone pool in the cathodic half-cell is oxidized (b = 0.1). The 80-150 mV range is only obtained for cells where more than 98% of the menaquinone pool in the anodic half-cell are reduced.

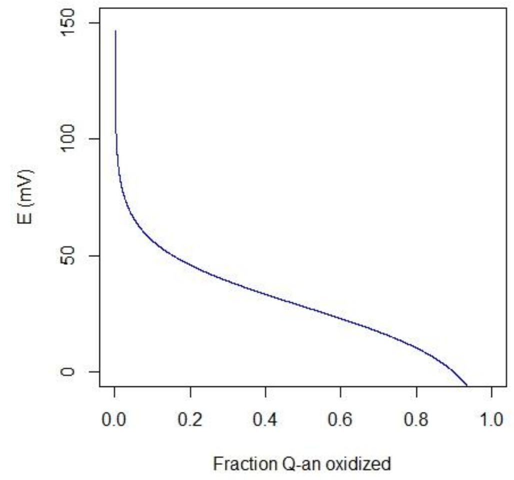


***Figure S5.3***. *The voltage yield of a concentraion cell as function of the fraction of menaquinons oxidized in the cathodic half-cell (b) where 90% of the menaquinone pool in the cathodic half-cell are oxidiced.*

### The current in the concentration cell

The above considerations points to the idea that sulfide oxidation coupled to nitrate reduction via menaquinone cycling, through long-distance electron transfer, is thermodynamically possible and the next question to be adressed is the if the voltage generated from the concentraion cell in a certain configuration can drive an electric current comparable to the current running in metabolic active cable bacteria. For simplicity, we will consider a steady state situation. The drivers of the model are the concentrations of oxidized and reduced menaquinones in the anodic and cathodic half-cells (eq. 1) We will assume that the redox state of the menaqunonens in the anodic and cathodic cells of the cable bacteria are controlled by 1) the rate of menaquinone reduction via sulfide oxidation in the anodic cells, 2)The rate of menaquinone oxidation in the anodic cells via the current generated from the concentration cell. 3) the rate of menaquinone reduction in the cathodic cells via the current generated from the concentration cell, and 3) the rate of menaquinone oxidation via nitrate reduction. At steady-state, (i.e. a, and b are constant) these rates are equal in magnitude. We will further assume that the total pool of menaquinones ($MQ_{Tot}$) in the anodic cell equals the total pool in the total pool of menaquinones ($MQ_{Tot}$) in the cathodic cells and that the rate constants menaquinone reduction in the cell is equal to the rate constant for menaquinone oxidation in the cathodic cell. This implies that a = (1-b) in equation S9, which then can be simplified to:

$$E = -\frac{0.0592}{2} Log\left(\frac{b^2 - 2b + 1}{b^2}\right) \qquad \text{(Eq. S10)}$$

With this expression we can estimate the curent in the steadystate sitution by means of ohms law.

I=E/R

Where I is the current, E the voltage yield of concentraion cell and R is the resistance of the wire.

The resistance of wire can be determined as in (3), *i.e.*,:

$$R = \frac{l}{\sigma A} \qquad \text{(Eq. S11)}$$

Here l is the length of the wire, σ the conductivity and A the cross section area of the wire.

The current can therfore be estimated as:

$$I = -\frac{0.0592}{2} Log\left(\frac{b^2 - 2b + 1}{b^2}\right)\frac{\sigma A}{l} \qquad \text{(Eq. S12)}$$

The conductive element in cable bacteria is a periplasmatic network of discrete fibres (4). Each of those has a diameter of ca. 50 nm, corresponding to a cross section area of $2 \times 10^{-15}$ $m^2$ and a counductivity of 20.1 S $cm^{-1}$ corresponding to $2.01 \times 10^3$ S $m^{-1}$ (4).

Figure S5.4, shows the current running between two half cells, connected with a wire having conductive properties similar to a cable bacteria fibre. The current flows from the anocic half cell the cathodic half cell if the fraction of oxidized menaquinones to the total pool of menaquinones in the cathodic half cell (b) is > 0.5. The current increases for b->1. There is an inverse relationship between the length of the wire connecting the anodic and cathodic half-cells and the magnitude of the current running through it. Half cells connected with a short wire in general has a higher current flowing between than half-cells connected with a long wire.

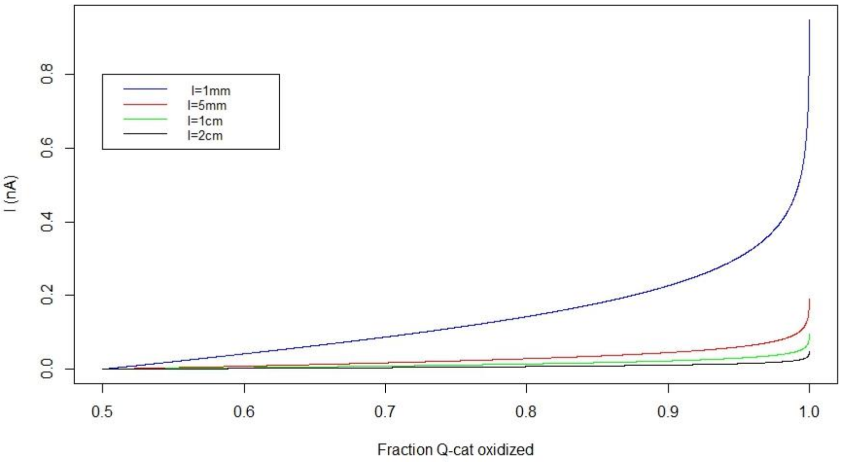


**Figure S5.4**. *Steady state current between half cells in the concentration cell, as function of b: the fraction of oxidized to total menaquinenes in the cathodic half cell. The half cells are electrically connected with a wire having conductive properties similar to a conducting fibre in cable bacteria. Collored lines represent the current in wires at different length (l).*

A cable bacterium can be considered as a composit of concentration cells with a series of anodic half cells coupled few coupled to a few cathodic half cells via the conducting periplasmatic network of fibres. We will for simplicity consider a virtual cable bacterium with 5000 anodic cells, each coupled to a cathodic cell. The cathodic cell can in principple be the same for all anodic cells. 5000 cells spans a distance of 2 cm and in our simplified model the wire connecting the most distant anodic cell to the cathodic cell is 2 cm long while the wire connecting the nearst anodic cell to the cathodic cell is 4 µm long. In general the length of the wire of the cells in between is equal to their distance to the cathodic cell, wich is in a fixed position for all anodic cells. According to equation S12 and Fig S5.4, the implication of such configuration is that the metabolic activity of the cells in the filament decreases with the distance to the cathodic cell. Figure S5.5 shows the current integrated for all 5000 cells in the filament. As seen the current generated from the this composit concentration cell model of a cable bacteria wastly exeed the 0.2 - 0.36 nA estimated for natural cable bacteria (5-7) for most redox states of the menaquinone pool in the cathodic cell. Already, with 50.11% of the menaquinones in the cathodic cell beeing oxidizede the curren is > 1 nA.

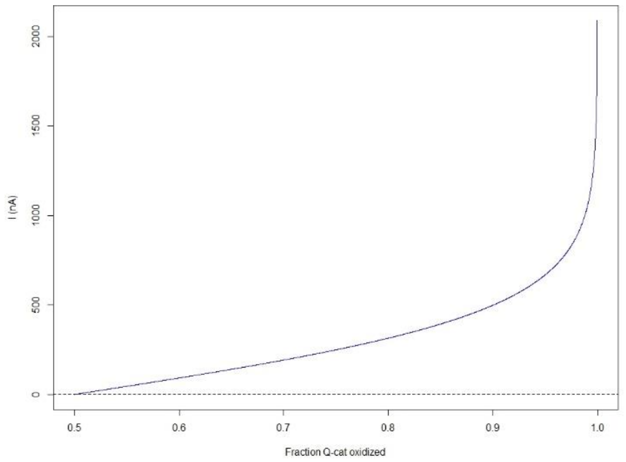


***Figure S5.5***. *Curent produced from 5000 individual half-cells (the app number of cells in a 2 cm long filament). Dashed line indicates the max filament specific current production (0.4 nA) reported in the literature.*

**Conclusion**

Here we have shown using the concentration cell model , that electron transport from a menaquinone pool in a anodic cell to a menaquinone pool in a cathodic cell is thermodynamically possible and that the voltage produced from such a cell can be sufficiently high to drive an electric current that exceeds the current reported for cable bacteria, when the concentration cells are brought together in a way that simulate a cable bacterium.
