## Supplementary material for "Dissimilatory Nitrate Reduction to Ammonium in the Cable Bacterium *Ca*. Electronema Sp. GS": Table S1

**Table S1: Transcript levels (RPKM) and differential transcription (DESeq2) of the 100 most highly expressed genes of *Ca* . Electronema sp. GS. DNRA genes are highlighted.**

| Rank | IMG Gene I | Locus Tag | Gene Product Name | Average RPKM under nitrate-reducing conditions | log2 fold Change between oxic control and nitrate amendment | Adjusted p value |
|---|---|---|---|---|---|---|
| 1 | 2729874347 | Ga0183576_10762 | PilA | 107488 | -1.12 | 1.00 |
| 2 | 2729875496 | Ga0183576_13321 | hypothetical protein | 37779 | 0.81 | 1.00 |
| 3 | 2729875974 | Ga0183576_16010 | cytochrome c | 36783 | 0.41 | 1.00 |
| 4 | 2729875937 | Ga0183576_1578 | hypothetical protein | 35366 | 1.52 | 0.96 |
| 5 | 2729874243 | Ga0183576_10656 | hypothetical protein | 29243 | -2.78 | 0.65 |
| 6 | 2729874606 | Ga0183576_1117 | hypothetical protein | 25514 | 0.34 | 1.00 |
| 7 | 2729874008 | Ga0183576_10443 | Putative peptidoglycan binding domain-containing protein | 19765 | 0.29 | 1.00 |
| 8 | 2729875690 | Ga0183576_14111 | hypothetical protein | 19659 | -0.11 | 1.00 |
| 9 | 2729875544 | Ga0183576_13513 | hypothetical protein | 16289 | -0.58 | 1.00 |
| 10 | 2729873391 | Ga0183576_10111 | cold-shock DNA-binding protein family | 16178 | 0.66 | 1.00 |
| 11 | 2729873390 | Ga0183576_10110 | cold-shock DNA-binding protein family | 13896 | -0.57 | 1.00 |
| 12 | 2729874498 | Ga0183576_10928 | hypothetical protein | 12838 | 0.37 | 1.00 |
| 13 | 2729875691 | Ga0183576_14112 | hypothetical protein | 12251 | -2.02 | 0.98 |
| 14 | 2729875791 | Ga0183576_14613 | hypothetical protein | 11553 | -0.14 | 1.00 |
| 15 | 2729874205 | Ga0183576_10618 | periplasmic nitrate reductase chaperone NapD | 11247 | 2.16 | 0.97 |
| 16 | 2729874203 | Ga0183576_10616 | periplasmic nitrate reductase subunit NapA apoprotein | 11063 | 1.09 | 1.00 |
| 17 | 2729875033 | Ga0183576_11921 | tRNA 2-thiouridine synthesizing protein E | 11022 | 0.95 | 1.00 |
| 18 | 2729874199 | Ga0183576_10612 | pMHC | 10387 | 0.76 | 1.00 |
| 19 | 2729874502 | Ga0183576_10932 | hypothetical protein | 10266 | 0.11 | 1.00 |
| 20 | 2729874460 | Ga0183576_10883 | hypothetical protein | 9917 | 1.68 | 0.76 |
| 21 | 2729874759 | Ga0183576_11353 | Opacity protein | 9686 | 0.78 | 1.00 |
| 22 | 2729873817 | Ga0183576_102205 | cold-shock DNA-binding protein family | 9540 | 1.78 | 0.76 |
| 23 | 2729874596 | Ga0183576_11052 | hypothetical protein | 9508 | -0.84 | 1.00 |
| 24 | 2729875792 | Ga0183576_14614 | 4Fe-4S dicluster domain-containing protein | 9457 | 0.35 | 1.00 |
| 25 | 2729874206 | Ga0183576_10619 | periplasmic nitrate reductase maturation protein NapF | 8988 | 1.83 | 1.00 |
| 26 | 2729875357 | Ga0183576_12812 | hypothetical protein | 8953 | -0.22 | 1.00 |
| 27 | 2729875626 | Ga0183576_1392 | pyrroloquinoline quinone biosynthesis protein D | 8725 | -0.74 | 1.00 |
| 28 | 2729874566 | Ga0183576_11022 | protein of unknown function (DUF4360) | 8432 | 1.85 | 0.89 |
| 29 | 2729873552 | Ga0183576_101173 | hypothetical protein | 8273 | -0.90 | 1.00 |
| 30 | 2729873752 | Ga0183576_102140 | LSU ribosomal protein L10P | 8041 | 0.86 | 1.00 |
| 31 | 2729874198 | Ga0183576_10611 | hypothetical protein | 7954 | -1.03 | 1.00 |
| 32 | 2729875893 | Ga0183576_1535 | hypothetical protein | 7912 | 0.13 | 1.00 |
| 33 | 2729873736 | Ga0183576_102124 | dissimilatory adenylylsulfate reductase beta subunit | 7383 | 0.04 | 1.00 |
| 34 | 2729873441 | Ga0183576_10161 | Dissimilatory sulfite reductase D (DsrD) | 7310 | 0.44 | 1.00 |
| 35 | 2729874204 | Ga0183576_10617 | periplasmic nitrate reductase chaperone NapD | 7118 | 2.22 | 0.98 |
| 36 | 2729873786 | Ga0183576_102174 | SSU ribosomal protein S4P | 7036 | 0.55 | 1.00 |
| 37 | 2729874424 | Ga0183576_10847 | integration host factor subunit beta | 6979 | 0.16 | 1.00 |
| 38 | 2729873757 | Ga0183576_102145 | SSU ribosomal protein S7P | 6792 | -0.04 | 1.00 |
| 39 | 2729874342 | Ga0183576_10757 | type IV pilus assembly protein PilW | 6786 | -0.03 | 1.00 |
| 40 | 2729874202 | Ga0183576_10615 | ferredoxin-type protein NapG | 6696 | 0.97 | 1.00 |
| 41 | 2729873774 | Ga0183576_102162 | small subunit ribosomal protein S14 | 6654 | 0.14 | 1.00 |
| 42 | 2729873784 | Ga0183576_102172 | small subunit ribosomal protein S13 | 6594 | -0.22 | 1.00 |
| 43 | 2729875627 | Ga0183576_1393 | hypothetical protein | 6565 | -0.52 | 1.00 |
| 44 | 2729876010 | Ga0183576_1637 | protein refolding chaperone Spy/CpxP family | 6469 | 1.51 | 1.00 |
| 45 | 2729875938 | Ga0183576_1579 | conserved repeat domain-containing protein | 6393 | 0.52 | 1.00 |
| 46 | 2729873766 | Ga0183576_102154 | LSU ribosomal protein L22P | 6281 | 0.40 | 1.00 |
| 47 | 2729873765 | Ga0183576_102153 | SSU ribosomal protein S19P | 6153 | -0.37 | 1.00 |
| 48 | 2729873773 | Ga0183576_102161 | LSU ribosomal protein L5P | 6105 | 0.11 | 1.00 |
| 49 | 2729873761 | Ga0183576_102149 | LSU ribosomal protein L3P | 5695 | 0.41 | 1.00 |
| 50 | 2729875790 | Ga0183576_14612 | hypothetical protein | 5618 | 0.95 | 1.00 |
| 51 | 2729875517 | Ga0183576_13418 | Chemoreceptor zinc-binding domain-containing protein | 5599 | -0.80 | 1.00 |
| 52 | 2729875936 | Ga0183576_1577 | hypothetical protein | 5557 | -1.23 | NA |
| 53 | 2729873737 | Ga0183576_102125 | dissimilatory adenylylsulfate reductase alpha subunit precursor | 5542 | 0.16 | 1.00 |
| 54 | 2729873628 | Ga0183576_10216 | hypothetical protein | 5520 | -1.23 | 1.00 |
| 55 | 2729873772 | Ga0183576_102160 | large subunit ribosomal protein L24 | 5453 | 0.26 | 1.00 |
| 56 | 2729873767 | Ga0183576_102155 | SSU ribosomal protein S3P | 5255 | 0.45 | 1.00 |
| 57 | 2729876009 | Ga0183576_1636 | protein of unknown function (DUF4405) | 5130 | 1.29 | 1.00 |
| 58 | 2729874874 | Ga0183576_1164 | hypothetical protein | 5022 | 0.12 | 1.00 |
| 59 | 2729875244 | Ga0183576_12431 | Protein of unknown function (DUF3106) | 4929 | -1.13 | 1.00 |
| 60 | 2729873775 | Ga0183576_102163 | SSU ribosomal protein S8P | 4912 | -0.12 | 1.00 |
| 61 | 2729873763 | Ga0183576_102151 | LSU ribosomal protein L23P | 4874 | 0.64 | 1.00 |
| 62 | 2729873994 | Ga0183576_10429 | hypothetical protein | 4783 | -0.83 | 1.00 |
| 63 | 2729874712 | Ga0183576_1136 | D-alanyl-D-alanine carboxypeptidase (penicillin-binding protein 5/6) | 4742 | 1.36 | 1.00 |
| 64 | 2729874102 | Ga0183576_10531 | transcriptional regulator, MucR family | 4710 | -0.20 | 1.00 |
| 65 | 2729874459 | Ga0183576_10882 | hypothetical protein | 4699 | 1.64 | 0.92 |
| 66 | 2729874605 | Ga0183576_1116 | hypothetical protein | 4621 | -0.40 | 1.00 |
| 67 | 2729874340 | Ga0183576_10755 | PilX N-terminal | 4597 | 0.89 | 1.00 |
| 68 | 2729875483 | Ga0183576_1338 | SSU ribosomal protein S21P | 4589 | 0.77 | 1.00 |
| 69 | 2729873785 | Ga0183576_102173 | SSU ribosomal protein S11P | 4513 | 0.62 | 1.00 |
| 70 | 2729874200 | Ga0183576_10613 | periplasmic nitrate reductase subunit NapB | 4490 | 0.45 | 1.00 |
| 71 | 2729875622 | Ga0183576_13819 | mRNA interferase YafQ | 4465 | -2.08 | 0.92 |
| 72 | 2729874323 | Ga0183576_10738 | hypothetical protein | 4441 | 0.31 | 1.00 |
| 73 | 2729873985 | Ga0183576_10420 | integration host factor subunit alpha | 4329 | -0.62 | 1.00 |
| 74 | 2729873758 | Ga0183576_102146 | translation elongation factor 2 (EF-2/EF-G) | 4263 | -0.01 | 1.00 |
| 75 | 2729875693 | Ga0183576_14114 | hypothetical protein | 4176 | -0.45 | 1.00 |
| 76 | 2729873987 | Ga0183576_10422 | hypothetical protein | 4138 | 1.08 | 1.00 |
| 77 | 2729873776 | Ga0183576_102164 | large subunit ribosomal protein L6 | 4109 | -0.07 | 1.00 |
| 78 | 2729873787 | Ga0183576_102175 | DNA-directed RNA polymerase subunit alpha | 3881 | 0.51 | 1.00 |
| 79 | 2729875722 | Ga0183576_14220 | manganese-dependent inorganic pyrophosphatase | 3820 | -0.13 | 1.00 |
| 80 | 2729873751 | Ga0183576_102139 | LSU ribosomal protein L1P | 3793 | 0.07 | 1.00 |
| 81 | 2729873778 | Ga0183576_102166 | SSU ribosomal protein S5P | 3776 | 0.13 | 1.00 |
| 82 | 2729874463 | Ga0183576_10886 | Uncharacterized conserved protein YkwD, contains CAP (CSP/antigen 5/PR1) domain | 3756 | -0.22 | 1.00 |
| 83 | 2729873745 | Ga0183576_102133 | translation elongation factor 1A (EF-1A/EF-Tu) | 3705 | 0.13 | 1.00 |
| 84 | 2729873993 | Ga0183576_10428 | hypothetical protein | 3643 | -1.36 | 0.98 |
| 85 | 2729875666 | Ga0183576_14017 | LSU ribosomal protein L13P | 3639 | 0.71 | 1.00 |
| 86 | 2729874304 | Ga0183576_10719 | hemoglobin | 3634 | 0.83 | 1.00 |
| 87 | 2729875208 | Ga0183576_12331 | Putative beta-barrel porin 2 | 3624 | -0.78 | 1.00 |
| 88 | 2729875284 | Ga0183576_12610 | chaperonin GroES | 3578 | 1.20 | 0.62 |
| 89 | 2729875916 | Ga0183576_1555 | outer membrane protein | 3530 | -1.55 | 1.00 |
| 90 | 2729875283 | Ga0183576_1269 | chaperonin GroEL | 3508 | 0.86 | 1.00 |
| 91 | 2729873762 | Ga0183576_102150 | LSU ribosomal protein L4P | 3507 | -0.08 | 1.00 |
| 92 | 2729873749 | Ga0183576_102137 | transcription antitermination protein nusG | 3503 | -0.22 | 1.00 |
| 93 | 2729873753 | Ga0183576_102141 | LSU ribosomal protein L12P | 3413 | 0.79 | 1.00 |
| 94 | 2729873764 | Ga0183576_102152 | LSU ribosomal protein L2P | 3403 | 0.39 | 1.00 |
| 95 | 2729875667 | Ga0183576_14018 | SSU ribosomal protein S9P | 3354 | 0.55 | 1.00 |
| 96 | 2729874201 | Ga0183576_10614 | ferredoxin-type protein NapH | 3235 | 1.88 | 0.96 |
| 97 | 2729873760 | Ga0183576_102148 | SSU ribosomal protein S10P | 3192 | 0.40 | 1.00 |

| | | | | | | |
|---|---|---|---|---|---|---|
| 98 | 2729874739 | Ga0183576_11333 | hypothetical protein | 3147 | 0.23 | 1.00 |
| 99 | 2729873783 | Ga0183576_102171 | bacterial translation initiation factor 1 (bIF-1) | 3113 | 0.65 | 1.00 |
| 100 | 2729874509 | Ga0183576_10939 | SSU ribosomal protein S1P | 3073 | 0.12 | 1.00 |

**Table S2: Accession numbers/gene identifiers for all sequences used for phylogenetic analysis of Nap genes (Fig. S1)**

| Species Name | Gene identifier [IMG/NCBI ID] |
|---|---|
| Candidatus Electronema sp. GS | 2729874203 |
| Aggregatibacter actinomycetemcomitans DSM 8324 | 2515250942 |
| Anaeromyxobacter dehalogenans 2CP-C | ABC80683.1 |
| Azospirillum brasilense sp7 | 2599104047 |
| Azospirillum brasilense sp7 | 2599101664 |
| Bordetella bronchiseptica RB50 | CAE33292.1 |
| Bordetella parapertussis 12822 | Q7W733.1 |
| Bradyrhizobium japonicum USDA 110 | 637374628 |
| Burkholderia xenovorans LB400 | 637953004 |
| Campylobacter jejuni jejuni NCTC 11168 | YP_002344187.1 |
| Candidatus Desulfuromonas soudanensis WTL | 2609285564 |
| Cupriavidus necator pHG1 | 640427642 |
| Cystobacter violaceus Cb vi76 | 2592593039 |
| Dechloromonas aromatica RCB | 637681987 |
| Desulfitobacterium hafniense DCB-2 | ACL19345.1 |
| Desulfobulbus japonicus DSM 18378 | WP_028579228.1 |
| Desulfobulbus mediterraneus | WP_028585221.1 |
| Desulfobulbus propionicus DSM 2032 | ADW17542.1 |
| Desulfocapsa sulfexigens DSM 10523 | AGF79815.1 |
| Desulfocapsa sulfexigens | WP_015405499.1 |
| Desulfovibrio desulfuricans ATCC 27774 | ACL48525.1 |
| Desulfurivibrio alkaliphilus AHT2 | 646847134 |
| Dissulfuribacter thermophilus S69 | OCC14250.1 |
| Escherichia coli DH1 | 646935360 |
| Geobacter lovleyi SZ | ACD94779.1 |
| Geobacter pickeringii G13 | AJE02263.1 |
| Haemophilus ducreyi ATCC 33940 | 2599173274 |
| Haemophilus influenzae F3047 | 649868883 |
| Helicobacter hepaticus ATCC 51449 | Q7VJT5.1 |
| Kryptonium sp. JGI-4 | 2599799712 |
| Kryptonium sp. JGI-6 | 2600397245 |
| Kryptonium sp. JGI-23 | 2601849902 |
| Magnetospirillum magneticum AMB-1 | 637821302 |
| Paracoccus denitrificans PD1222 | ABL72782.1 |
| Paracoccus pantotrophus DSM 1403 | SFY43853.1 |
| Pasteurella multocida 36950 | 2512391891 |
| Photobacterium profundum SS9 | 637585562 |
| Photobacterium profundum SS9 | 637586736 |
| Pseudomonas aeruginosa PAO1 | 637051567 |
| Pseudomonas sp. G-179 | AAD46689.1 |
| Ralstonia eutropha JMP134 | 637694190 |
| Ralstonia metallidurans CH34 | 637979487 |
| Rhizobium sp. CF142 | WP_007818974.1 |
| Rhodobacter sphaeroides 2.4.1 | ABA81591.1 |
| Salmonella enterica enterica sv. Choleraesuis SC-B67 | 637641152 |
| Salmonella typhimurium LT2 | 637212978 |
| Sedimenticola selenatireducens AK4OH1 | 2513982609 |
| Sinorhizobium meliloti | AIM03004.1 |
| Shewanella frigidimarina NCIMB 400 | ABI70414.1 |
| Shewanella oneidensis MR-1 | AAN53924.1 |
| Shigella flexneri CCH060 | 2531700739 |
| Sinorhizobium meliloti RMO17 | 2598504540 |
| Sorangium cellulosum 'So ce 56' | 641348992 |
| Sulfurospirillum barnesii SES-3 | 2507135060 |
| Symbiobacterium thermophilum IAM 14863 | BAD39902.1 |
| Thermosulfurimonas dismutans | OAQ21381.1 |
| Vibrio vulnificus ATCC 43382 | 2662388500 |
| Wolinella succinogenes DSM 1740 | 637456405 |
| Bacillus subtilis | WP_124073427.1 |
| Synechococcus elongatus PCC 7942 | CAA52675.1 |

**Table S3: Accession numbers/gene identifiers for all sequences used for phylogenetic analysis of NapD genes (Fig. S2)**

| | |
|---|---|
| Candidatus Electronema sp. GS | 2729874204 |
| Candidatus Electronema sp. GS | 2729874205 |
| Desulfurivibrio alkaliphilus AHT2 | 646847137 |
| Aggregatibacter actinomycetemcomitans DSM 8324 | 2515250943 |
| Anaeromyxobacter dehalogenans 2CP-C | ABC80682.1 |
| Azospirillum brasilense sp7 | 2599104048 |
| Azospirillum brasilense sp7 | 2599101665 |
| Bordetella bronchiseptica RB50 | CAE33291.1 |
| Bordetella parapertussis 12822 | 637109774 |
| Bradyrhizobium japonicum USDA 110 | 637374627 |
| Burkholderia xenovorans LB400 | 637953003 |
| Campylobacter jejuni jejuni NCTC 11168 | CAL34913.1 |
| Candidatus Desulfuromonas soudanensis WTL | 2609285565 |
| Cupriavidus necator pHG1 | 640427641 |
| Dechloromonas aromatica RCB | 637681988 |
| Desulfitobacterium hafniense DCB-2 | 643561198 |
| Desulfobulbus japonicus DSM 18378 | WP_028579227.1 |
| Desulfobulbus propionicus DSM 2032 | ADW17543.1 |
| Desulfocapsa sulfexigens DSM 10523 | AGF78837.1 |
| Desulfovibrio desulfuricans ATCC 27774 | ACL48526.1 |
| Desulfurivibrio alkaliphilus AHT2 | 646847137 |
| Escherichia coli DH1 | 646935359 |
| Geobacter lovleyi SZ | ACD94778.1 |
| Geobacter pickeringii G13 | AJE02262.1 |
| Haemophilus ducreyi ATCC 33940 | 2599173275 |
| Haemophilus influenzae F3047 | 649868884 |
| Helicobacter hepaticus ATCC 51449 | 637431176 |
| Kryptonium sp. JGI-4 | 2599799715 |
| Kryptonium sp. JGI-6 | 2600397248 |
| Magnetospirillum magneticum AMB-1 | BAE51495.1 |
| Paracoccus denitrificans PD1222 | ABL72781.1 |
| Paracoccus pantotrophus | SFY43855.1 |
| Pasteurella multocida 36950 | 2512391890 |
| Photobacterium profundum SS9 | 637585561 |
| Pseudomonas aeruginosa PAO1 | 637051568 |
| Pseudomonas sp. G-179 | AAD46688.1 |
| Ralstonia metallidurans CH34 | 637979488 |
| Rhizobium sp. CF142 | EJJ28403.1 |
| Rhodobacter sphaeroides 2.4.1 | 640069464 |
| Salmonella enterica enterica sv. Choleraesuis SC-B67 | 637641153 |
| Salmonella typhimurium LT2 | 637212979 |
| Sedimenticola selenatireducens AK4OH1 | 2513982608 |
| Shewanella frigidimarina NCIMB 400 | ABI72396.1 |
| Shewanella oneidensis MR-1 | AAN53925.1 |
| Shigella flexneri CCH060 | 2531700738 |
| Sinorhizobium meliloti RMO17 | 2598504539 |
| Sulfurospirillum barnesii SES-3 | 2507135066 |
| Symbiobacterium thermophilum IAM 14863 | 637537430 |
| Wolinella succinogenes DSM 1740 | 637456399 |

**Table S4: Accession numbers/gene identifiers for all sequences used for phylogenetic analysis of NapF genes (Fig. S3)**

| | |
|---|---|
| Candidatus Electronema sp. GS | 2729874206 |
| Candidatus Electrothrix aarhusiensis MCF | 2608444788 |
| Candidatus Electrothrix marina A3 | 2609183527 |
| Aggregatibacter segnis | WP_109859830.1 |
| Arcobacter sp. CECT 8986 | WP_128990985.1 |
| Avibacterium avium | WP_115250139.1 |
| Azospirillum brasilense sp7 | 2599104049 |
| Azovibrio restrictus | WP_026685580.1 |
| Beggiatoa sp. 4572 | OQY57412.1 |
| Candidatus Accumulibacter sp. | TLD45631.1 |
| Denitromonas halophilus | WP_144176374.1 |
| Enterovibrio coralii | WP_067418395.1 |
| Escherichia coli DH1 | 646935358 |
| Haemophilus ducreyi ATCC 33940 | 2599173276 |
| Haemophilus influenzae F3047 | 649868885 |
| Hydrogenimonas thermophila | WP_092910301.1 |
| Magnetospira sp. QH-2 | WP_046020781.1 |
| Magnetospirillum magneticum AMB-1 | 637821304 |
| Pasteurella multocida 36950 | 2512391889 |
| Pectobacterium peruviense | WP_048259758.1 |
| Pectobacterium zantedeschiae | WP_129706134.1 |
| Photobacterium profundum SS9 | 637585560 |
| Pseudomonas aeruginosa PAO1 | 637051569 |
| Pseudomonas sp. G-179 | AAD46687.1 |
| Rhizobium sp. CF142 | WP_007818978.1 |
| Rhodocyclaceae bacterium | TXG88186.1 |
| Rodentibacter ratti | WP_077497317.1 |
| Salmonella typhimurium LT2 | 637212980 |
| Sedimenticola selenatireducens AK4OH1 | 2513982607 |
| Shewanella frigidimarina NCIMB 400 | ABI71179.1 |
| Shewanella oneidensis MR-1 | AAN54718.1 |
| Shigella flexneri CCH060 | 2531700737 |
| Sinorhizobium meliloti RMO17 | 2598504538 |
| Solemya elarraichensis gill symbiont | WP_078476978.1 |
| Sulfuricella denitrificans | WP_148290815.1 |
| Sulfurospirillum barnesii SES-3 | 2507135064 |
| Vibrio vulnificus ATCC 43382 | 2662388498 |
| Wolinella succinogenes DSM 1740 | 637456401 |

**Table S5: Accession numbers/gene identifiers for all sequences used for phylogenetic analysis of pMHC/εHao/NrfA genes (Fig. S4)**

| | |
|---|---|
| Candidatus Electronema sp. GS | 2729874199 |
| Candidatus Electrothrix aarhusiensis MCF | 2608444786 |
| Beggiatoa sp. "Orange Guaymas" | 2502838749 |
| Candidatus Thiomargarita nelsonii S10 | 2601778522 |
| Candidatus Thioglobus autotrophicus | WP_053951924.1 |
| Caminibacter mediatlanticus | WP_007473950.1 |
| Campylobacter curvus | WP_011992073.1 |
| Campylobacter fetus | WP_002849252.1 |
| Nautilia profundicola | WP_015902282.1 |
| Actinobacillus pleuropneumoniae S8 | 2552278133 |
| Actinobacillus succinogenes 130Z | 640807619 |
| Actinobacillus ureae ATCC 25976 | 650340686 |
| Ammonifex degensii KC4 | 646359879 |
| Anaerococcus prevotii ACS-065-V-Col13 | 2529738369 |
| Anaerolinea thermophila UNI-1 | 649906650 |
| Bacillus sp. 1NLA3E | 2506744287 |
| Caldilinea aerophila STL-6-01, DSM 14535 | 2513225030 |
| Campylobacter hominis ATCC BAA-381 | 640869848 |
| Campylobacter jejuni jejuni NCTC 11168 | 637040891 |
| Capnocytophaga gingivalis JCVIHMP016 | 644450706 |
| Citrobacter koseri ATCC BAA-895 | 640917778 |
| Comamonadaceae bacterium EBPR | 2619975620 |
| Corynebacterium pseudotuberculosis CCUG 27541 | 2628297476 |
| Delta proteobacterium MLMS-1 | 639155534 |
| Desulfosporosinus meridiei S10 | 2510242216 |
| Desulfovibrio desulfuricans ATCC 27774 | 643580866 |
| Desulfovibrio salexigens DSM 2638 | 644838886 |
| Desulfovibrio vulgaris Hildenborough | 637121843 |
| Desulfurispirillum indicum S5 | 649844489 |
| Desulfurivibrio alkaliphilus AHT2 | 646845286 |
| Escherichia coli HS | 640923413 |
| Gallibacterium anatis DSM 16844 | 2514917813 |
| Geobacter uraniumreducens Rf4 | 640551483 |
| Haemophilus influenzae 2019 | 2630849089 |
| Marivirga tractuosa DSM 4126 | 649787403 |
| Meiothermus silvanus DSM 9946 | 646843608 |
| Myxococcus xanthus DK 1622 | 638023578 |
| Opitutus terrae PB90-1 | 641694276 |
| Pelobacter carbinolicus DSM 2380 | 637752476 |
| Photobacterium profundum SS9 | 637585967 |
| Planctomyces brasiliensis DSM 5305 | 649980400 |
| Planctomyces maris DSM 8797 | 641112668 |
| Porphyromonas gingivalis W83 | 637150428 |
| Prevotella marshii DSM 16973 | 648809557 |
| Propionivibrio dicarboxylicus DSM 5885 | 2599430875 |
| Providencia alcalifaciens DSM 30120 | 643148102 |
| Riemerella anatipestifer DSM 15868 | 649778058 |
| Selenomonas sputigena ATCC 35185 | 646081185 |
| Shewanella oneidensis MR-1 | 637345732 |
| Sulfurospirillum deleyianum DSM 6946 | 646423959 |
| Thioalkalivibrio nitratireducens DSM 14787 | 2521963319 |
| Thioalkalivibrio paradoxus ARh 1 | 2513007479 |
| Thioalkalivibrio thiocyanoxidans ARh 4 | 2506728454 |
| Vibrio fischeri ES114 | 637636232 |
| Vibrio harveyi ATCC 25919 | 2583678091 |
| Wolinella succinogenes DSM 1740 | 637456212 |
| Yersinia enterocolitica YEA | 2611555138 |
| Aeromonas hydrophila ATCC 7966 | 639715039 |
| Bacillus subtilis | KOS70037.1 |
